# The Hitchhiker’s Guide to the Periplasm: Unexpected Molecular Interactions of Antibiotics Revealed by Considering Crowding Effects in *E. coli*

**DOI:** 10.1101/2020.06.03.132118

**Authors:** Conrado Pedebos, Iain P. S. Smith, Alister Boags, Syma Khalid

## Abstract

The periplasm of Gram-negative bacteria is a highly crowded environment comprised of many different molecular species. Antibacterial agents that causes lysis of Gram-negative bacteria by their action against the inner membrane must cross the periplasm to arrive at their target membrane. Very little is currently known about their route through the periplasm, and the interactions they experience. To this end, here atomistic molecular dynamics simulations are used to study the path taken by the antibiotic polymyxin B1 through a number of models of the periplasm which are crowded with proteins and osmolytes to different extents. The simulations reveal that PMB1 forms transient and long-lived interactions with proteins and osmolytes that are free in solution as well as lipoproteins anchored to the outer membrane and bound to the cell wall. We show that PMB1 may be able to ‘hitchhike’ within the periplasm by binding to lipoprotein carriers. Overall our results show that PMB1 is rarely uncomplexed within the periplasm; an important consideration for interpretations of its therapeutic mechanism of action. It is likely that this observation can be extended to other antibiotics that rely on diffusion to cross the periplasm.

## Introduction

The periplasm of Gram-negative bacteria is a crowded aqueous compartment bounded by the inner and outer membranes. The cell wall is contained within the periplasm as well as hundreds of proteins including chaperones, transporters proteases and nucleases^1,2^. The solution also contains a range of osmolytes, including urea, sugars, spermidine and putrescine. This makes for a complex and crowded environment for any molecular species to negotiate when moving across the periplasm towards either membrane.

Very little is known about the spatial arrangement of these myriad molecules within the periplasm. In other words, it is not known if the proteins and osmolytes are evenly distributed, or if is there some degree of organization and if so, to what extent. This makes it very difficult to predict the interactions experienced by molecules within the periplasm. This extends to molecules that are not native to the bacteria, such as antibacterial agents. Thus, we have little information regarding which moieties of antibiotics are available to carry out the desired functions, and which are unavailable as they are involved in interactions with native proteins/osmolytes/cell wall. To this end we have conducted a study of polymyxin B1 (PMB1) within models of the *E. coli* periplasm. PMB1 is a lipopeptide antibiotic used as a “last resort” drug for the treatment of infections caused by Gram-negative bacteria^3^. PMB is composed of a cyclic, cationic polypeptide ring connected to a branched fatty acid tail. The cationic ring contains five residues of the irregular amino acid α,γ-Diamino Butyric acid (DAB), each of which contributes a charge of +1 *e* giving PMB1 an overall charge of +5 *e*. The cationic ring enables solubility in aqueous solvents, whereas the lipid tail facilitates insertion into bacterial membranes^4-7^. While PMB1 along with colistin (polymyxin E) were for many years, last resort antibiotics, in recent years bacterial strains that are resistant to both antibiotics have emerged in a number of countries^8,9^. Thus, in order to either modify these drugs or to develop completely novel antibiotics, it is timely to establish a thorough, molecular-level understanding of each stage of the process *via* which they bring about bacterial cell death. To date, mechanistic studies of PMB1 have focused almost entirely on the two membranes of Gram-negative bacteria^7,10^, leaving unaddressed the question, how does PMB1 cross the periplasm to get from the outer membrane to the inner membrane? Here, a series of atomistic molecular dynamics simulations (Table 1) were performed of models of portions of the *E. coli* cell envelope. The simulation systems contain an asymmetric model of the outer membrane composed of LPS and phospholipids, a single-layered cell wall, various proteins/lipoproteins, osmolytes and PMB1, with systems sizes ranging from 200,000 to 750,000 atoms. The proteins are a combination of Braun’s lipoprotein (BLP), LolA, LolB, OmpA, and Pal (Fig1). BLP is the most abundant protein in *E. coli* (there are an estimated 10^5^ copies of BLP in each *E. coli*^11^). It exists as a coiled-coil trimer that is essential for compartment stability^12^. It is anchored in the outer membrane *via* a lipidated moiety at its N-terminus, whereas it is covalently bound to peptidoglycan via its C-terminus. LolA and LolB are small soluble proteins that carry lipoproteins^13,14^, they are largely similar in structure, although LolB is anchored to the OM *via* a lipidated moiety whereas LolA is free to diffuse across the cell envelope. OmpA is composed of an eight-stranded barrel which is located in the OM, and is connected *via* a linker to the soluble domain that can bind peptidoglycan in the periplasm^15,16,17^. Pal also has a lipidated anchor in the OM like LolB, while its C-terminal domain resembles the OmpA soluble domain. Like OmpA, Pal also has a linker that can extend into the periplasm enabling the protein to bind non-covalently to peptidoglycan, thereby assisting with maintaining compartment integrity^18,19^. Where present, in each system there are 4 x BLPs and 1 x each of OmpA, Pal, LolA and LolB. The most compositionally complex system studied here also contained a range of osmolytes, in order to better represent the crowded environment that these molecules encounter in the periplasm.

**Figure 1.**
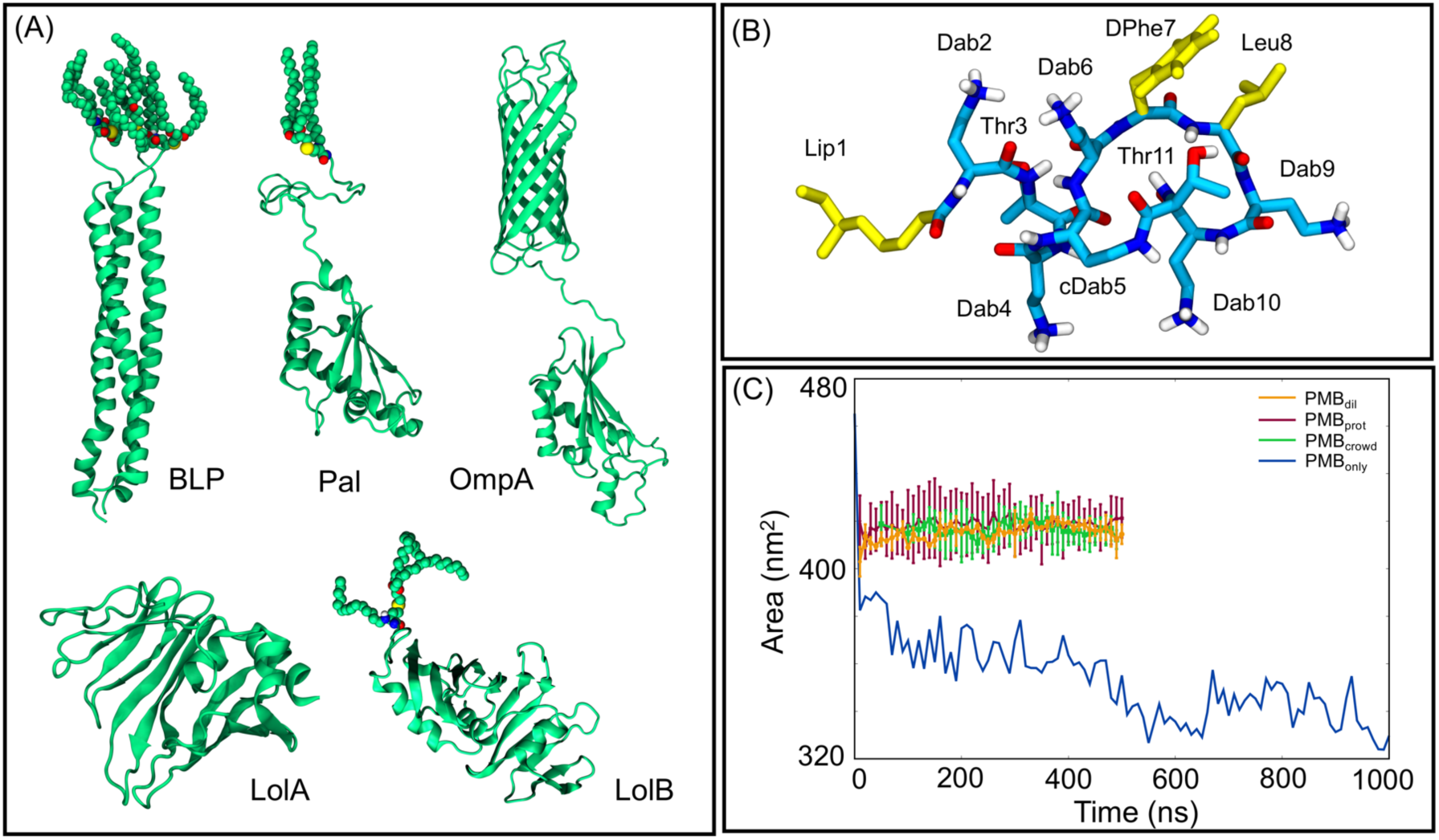
Summary of proteins studied and SASA data. Panel (A) shows the structures of the five proteins simulated in this study. Panel (B) shows the structure of PMB1 and panel (C) provides a summary of SASA versus time for each system.

**Table 1.** Summary of all simulated systems.

| Simulations |  |  |  |  |  |
| --- | --- | --- | --- | --- | --- |
| System | Proteins | PMB1 Molecules | Osmolytes and Ions<sup>a</sup> | Length | Force field |
| PMB<sub>only</sub> | None | 30 | Na<sup>+</sup>, Cl<sup>-</sup> | 1 μs | GROMOS 54a7 |
| PMB<sub>dil</sub> | BLP (x4) | 30 | Na<sup>+</sup>, Cl<sup>-</sup> | 2 x 500 ns | GROMOS 54a7 |
| PMB<sub>prot</sub> | BLP (x4), LolA, LolB, Pal, OmpA | 30 | Na<sup>+</sup>, Cl<sup>-</sup> | 2 x 500ns | GROMOS 54a7 |
| PMB<sub>crowd</sub> | BLP (x4), LolA, LolB, Pal, OmpA | 30 | Na<sup>+</sup>, Cl<sup>-</sup>, Glycerol, Urea, Trehalose, Spermidine, Putrescine, OPG | 2 x 500 ns | GROMOS 54a7 |

<sup>a</sup> Molar concentration: Na<sup>+</sup>, Cl<sup>-</sup> (200 mM), Glycerol (35mM), Urea (30mM), Trehalose (10 mM), Spermidine (0.2 mM), Putrescine (30 mM), Osmoregulated Periplasmic Glucans (OPG - 20 mM).

The osmolytes incorporated into our periplasmic model were selected on the basis of a combination of their abundance and chemical diversity. Importantly, all of these osmolytes have their concentrations in the periplasm either documented or estimated in the literature^20-25^ and these concentrations are reproduced here: osmoregulated periplasmic glucans (OPG) (20 mM), trehalose (10 mM), putrescine (30 mM), spermidine (3 mM), glycerol (36 mM) and urea (20 mM). Both OPG and trehalose are widely distributed in Bacteria, with OPG having a prominent role on regulating osmotic pressure and virulence^26^, whereas trehalose is mainly involved in response to stress conditions^27^. The polyamines, putrescine and spermidine, are the two most common in all bacteria, with functions that includes supporting bacterial growth, incorporation into the cell wall, and biosynthesis of siderophores^28^. Glycerol is metabolized in *E. coli* cells for different applications, both aerobically and anaerobically^29,30^. Urea is a source of nitrogen, after its breakdown^31^.

Simulations were initiated by placing PMB1 molecules randomly in the aqueous region between the outer membrane and the cell wall. The osmolyte concentrations are derived from literature values and the number of proteins is selected to reproduce crowding volume fraction of ϕ ∼0.21 as estimated from experimental studies^20^. A set of simulations of PMB1 in just water and ions was also performed for comparison.

## Results

Table 1 provided a summary of the simulations performed in this study. Initial observations focused on general mobility and aggregation of PMB1 followed by in depth analyses probing the causes of these observations.

The crowded nature of the systems had a clear impact upon the solvent accessible surface area (SASA) of PMB1 (Fig. 1). The SASA is lower when PMB1 molecules are just in water and counter ions (PMB1_only_), compared to when in the protein-containing systems (PMB1_dil_, PMB1_prot_ and PMB1_crowd_). Tracking the PMB1 motion within the XY plane (Fig. S1) of the protein-containing systems shows the movement of polymyxins in the crowded systems (PMB1_crowd_ and PMB1_prot_) is more confined compared to PMB1_dil_ in which BLP is the only protein. Additionally, in the latter system, more PMB1 molecules moved towards the outer membrane and the cell wall rather than remaining in the solution area between these two large structures, compared to PMB1_crowd_ and PMB1_prot_. Another effect observed with increasing system complexity is the slower diffusion of PMB1 (Fig. 2A and Fig. S2), by calculation of the translational diffusion coefficients (D_t_) from two different time regimes (1-10ns; and 50-100ns). For the PMB_only_ system, the D_t_ from the longer time regime was estimated to be 4.0 ± 0.3 × 10^−6^ cm^2^/s, while for PMB_dil_, PMB_prot,_ and PMB_crowd_ systems, the values were 3.8 ± 0.4 × 10^−8^ cm^2^/s, 2.9 ± 0.3 × 10^−8^ cm^2^/s and 2.0 ± 0.4 × 10^−8^ cm^2^/s, respectively, demonstrating a major reduction when compared to PMB_only_ (100-fold). The slowest diffusion is recorded for the most crowded system.

**Figure 2:**
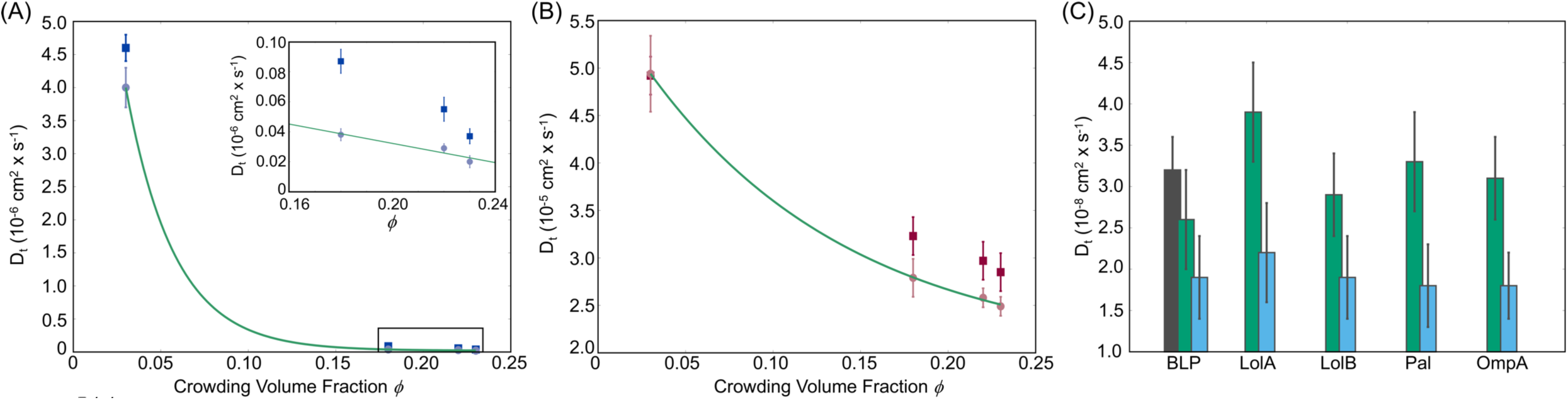
Translational diffusion coefficients (D_t_) obtained for PMB1 (left panel), water (middle panel), and proteins (right panel). (A): Dt values calculated for two different time regimes, 1-10 ns (blue) and 10-100 ns (grey), as a function of the crowding volume fraction ϕ of each system (PMB_only_ = 0.03; PMB_dil_ = 0.18; PMB_prot_ = 0.22; PMB_crowd_ = 0.23). (B): D_t_ values obtained for water molecules in the same time regimes as above, 1-10 ns (red) and 10-100 ns (pink), as a function of the crowding volume fraction ϕ. Exponential fits were applied for the long-time scale regimes. (C): Histogram showing D_t_ values for each protein in each system (PMB_dil_ = black; PMB_prot_ = green; PMB_crowd_ = blue). Error bars indicate standard error for all molecules across all repeat simulations.

The dynamics of water was also impacted by crowding (Fig. 2B), with a D_t_ rate of 4.93 ± 0.4 × 10^−5^ cm^2^/s in PMB_only_ compared to 2.49 ± 0.1 × 10^−5^ cm^2^/s in PMB_crowd_ for the 50-100 ns time period. The values found for systems in the presence of the outer membrane are similar to a previous report^32^ of simulations of the outer membrane in water using the SPC water model^33^ and with crowded simulations^34^ using a different water model. Protein diffusion rates were also calculated for the PMB_dil_, PMB_prot_, and PMB_crowd_ systems, showing D_t_ values that also decrease with increasing crowding volume fraction ϕ. Although LolA is neither bound to the cell wall nor anchored/embedded in the membrane, its calculated D_t_ falls in the same range as the other proteins indicating that overall protein motion is quite restricted in the crowded systems for all proteins. While the environment we have simulated is more complex due to the presence of membrane and cell wall, the diffusion rates for proteins calculated here are comparable to previous reports involving simulations of crowded environments^35^ and cytoplasm models^36,37^, as well as with experimental data from GFP proteins at the periplasm and cytoplasm^38^.

In this study we seek to characterize the molecular interactions that underpin the aforementioned SASA, lateral motion and translational diffusion profiles calculated from our simulations. The complexity of the system composition is such that a vast amount of data regarding molecular interactions is generated from these simulations. To facilitate interpretation of the observations we have presented the results from the perspective of PMB1 interactions, namely PMB1 interactions with itself, osmolytes, proteins, and the cell wall.

### PMB1 self-interactions

In systems containing only PMB1 in solution (PMB_only_), differently sized aggregates (dimers to pentamers) formed during the simulations. The lifetimes of interactions between PMB1 molecules ranged from short periods (a few nanoseconds) to longer term interactions (200-400 ns) leading to formation of aggregates, as shown in the example in Fig. 3A-B. A range of different configurations were observed during the simulations. The majority of the interactions occurred *via* the hydrophobic portions of PMB1, namely Lip1, DPhe7, and Leu8, while the charged sites remained largely exposed to water and ions (Fig. 3A-B). In the example of a tetrameric association as shown in Fig. 3A, four of the PMB1 molecules had their Lip1 tails buried in the middle of the aggregate along with two DPhe7 and three Leu8 moieties, thus forming a structure that resembled a micelle. Due to the exposure to the aqueous environment of the positive charges and polar residues in this tetramer, the surface of the micelle-like structure was decorated by Cl^-^ ions, which interacted mostly with the NH_3_^+^ groups from Dab residues. The center of the micelle was mostly protected from exposure to water (Fig. 3D), with only one constant water molecule present at 0.5 nm (Fig. S3). This self-assembly behavior has previously been reported for other similar amphiphilic antibiotics, such as colistin and colistin methanesulfonate (CMS), but shown not to occur for the non-amphiphilic polymyxin B nonapeptide, an analogue that lacks the hydrophobic tail^39^. In the cases previously reported, aggregate diameters were calculated to have a z-average of around 2 nm ± 0.3, which correlates well with the tetrameric aggregate observed in our simulations (2.2 nm ± 0.5). Thus, as predicted for colistin and its analogue^39^, PMB1 micelle formation followed a “closed association” model, where the number of monomers per micelle does not exceed five in our simulations. In the PMB_dil_, PMB_prot_, and PMB_crowd_ systems, interactions between PMB1s resulted in smaller aggregates, generally involving dimerization (but with the additional participation of other molecular species, as discussed in the next section).

**Figure 3:**
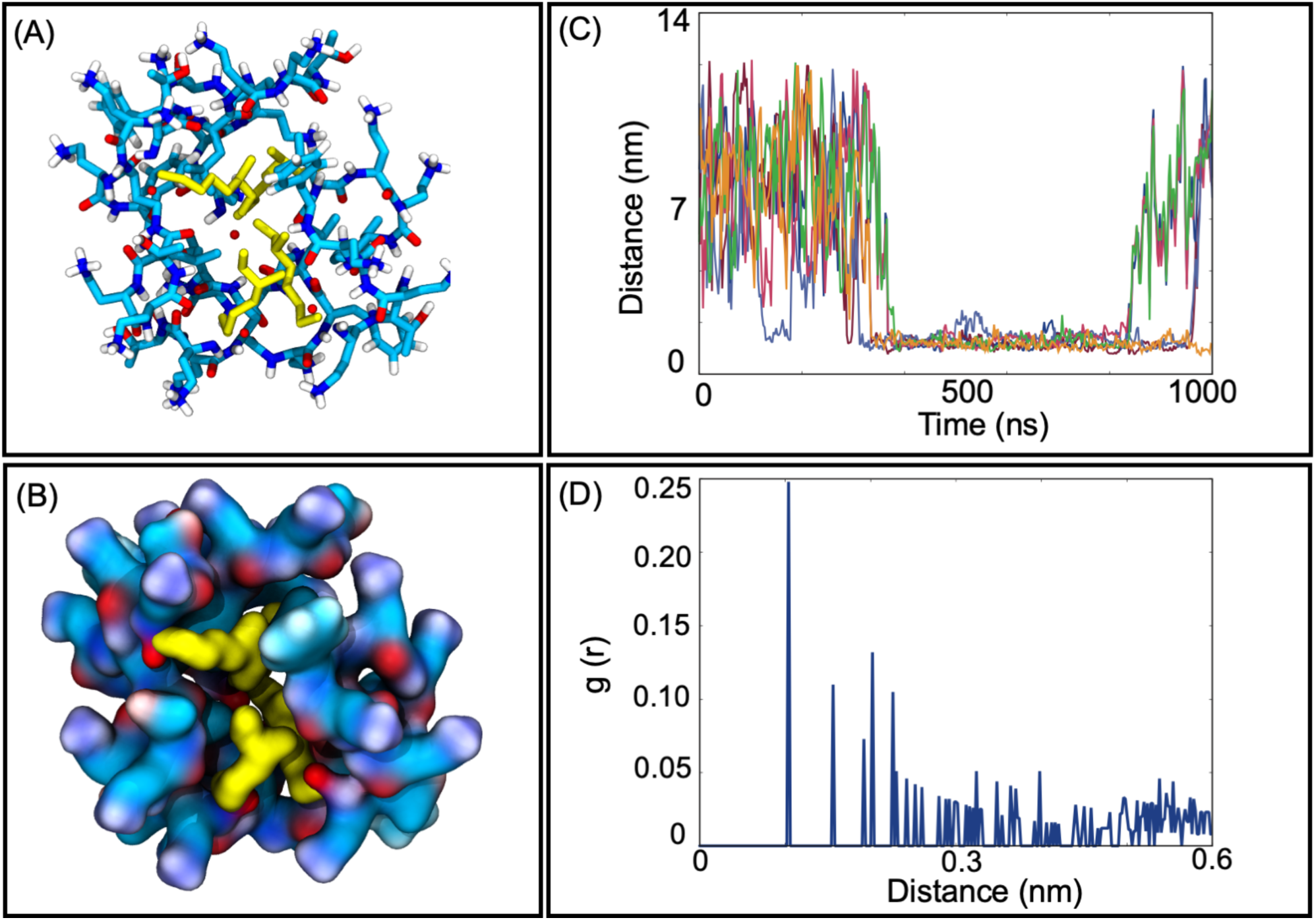
Micelle-like associations observed during the simulations. Sticks (A) and surface (B) representations of a tetrameric micelle, with the hydrophobic portions pointing inwards the aggregate colored in yellow. (C) Distances calculated between the center of mass (COM) of each monomer composing the micelle structure. Each curve describes the distance between two different monomers, with values below the 2 nm threshold indicating an association. (D) Radial Distribution Function (RDF) for water molecules calculated using the COM of the whole aggregate as a reference point.

### PMB1 interaction with osmolytes

The interaction of PMB1 with osmolytes and ions was firstly characterized by measuring the proximity of each type of osmolyte to PMB1 molecules. The radial distribution function (RDF) of each osmolyte with PMB1s as a reference (Fig. 4) showed a clear preference for glycerol and OPG. This is reasonable considering the number of polar groups available for interactions on both osmolytes and the negative charge (−1 *e*) on the phosphate group of OPGs. Putrescine, spermidine and Na^+^ ions were found furthest from PMB1, which correlates with both being positively charged (putrescine = +2 *e*, spermidine = +4 *e*).

**Figure 4:**
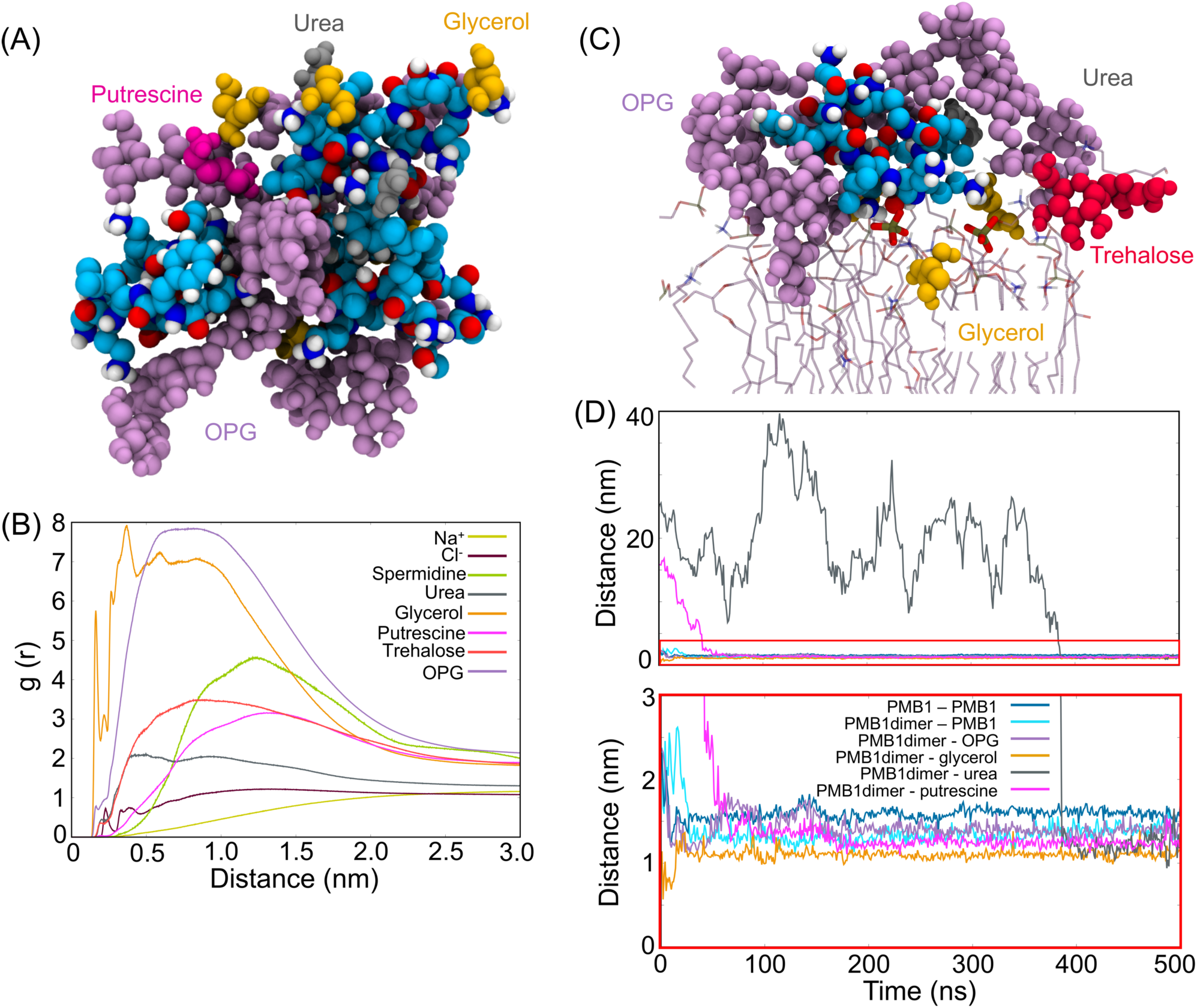
Osmolyte distribution and cluster formation in PMB_crowd_. (A) Cluster formed by three PMB1 molecules (cyan, white, red, blue), five OPG molecules (violet), four glycerol molecules (orange), two urea molecules (grey) and one putrescine molecule (magenta). (B) Radial distribution function (RDF) using PMB1 as a reference point with Glycerol (orange), OPG (violet) and Cl^-^ (maroon), Na^+^ (yellow), Putrescine (magenta) and Spermidine (light green), urea (grey) and trehalose (red). (C) Cluster formed at the surface of the outer membrane involving one PMB1 molecule, three OPG molecule, two glycerol molecules, one urea molecule and one trehalose molecule (colors as in (A)). Phosphate groups (brown and red sticks) form salt bridge interactions with Dab6 and Dab10 residues of PMB1. (D) Distances between representative molecules forming clusters are shown in panel (A), with a zoomed-in area marked with a red rectangle. Colored curves correspond to PMB1-PMB1, PMB1dimer-PMB1, PMB1dimer-OPG, PMB1dimer-glycerol and PMB1dimer-urea.

It has been discussed previously^35,40-44^ that in crowded cellular environments, non-specific binding occurs constantly, generating transient clusters that affect the structure and dynamics of the molecules in this environment. In the simulations performed here, we observed formation of small osmolyte-PMB1 clusters which had an average size ∼2.5 - 3.0 nm (slightly larger than the PMB1 micelles described in the previous section). These clusters generally contained PMB1 monomers interacting directly with OPG (*via* -OH groups and cyclohexane rings) and glycerol (*via* -OH groups), although participation of other osmolytes such as putrescine and urea was also observed, but usually without these molecules directly interacting with PMB1. In particular, the association between PMB1 molecules and OPG was prevalent (as shown in the RDF in Fig 4). For example, in one case, four OPG molecules bound around the surface of a PMB1 dimer (Fig. 4A), while a fifth OPG molecule mediated the interaction between the PMB1 dimer and a third PMB1 molecule. Four additional molecules of glycerol, one putrescine and two urea molecules also participated in this cluster, effectively bridging the PMB1 dimer to the third PMB1 (Fig. 4D), stabilizing the complex. This cluster took ∼ 100 ns to stabilize in terms of number of components, apart from one urea molecule which only joined the cluster after 400 ns (Fig. 4D and 4E). The final cluster shape was achieved at around 420 ns and remained stable until the end of the simulation. The largest cluster in all simulations was ∼ 4.2 nm in diameter and was composed of four PMB1 molecules and ∼20 osmolytes (one trehalose, five putrescine, seven glycerol and seven OPG). In this cluster, only two of the PMB1 molecules are directly associated with each other, interacting *via* their DPhe7 residues. The formation of the cluster was initiated by many of the molecules binding to the cell wall (within 30 ns of the start of the simulation), while the full cluster had formed after ∼100 ns and lasted for around 240 ns. Despite showing a higher preference for cluster formation in the cell wall area, a few aggregates were also observed on the surface of the inner leaflet of the outer membrane (Figure 4C). For example, in one cluster, one PMB1 molecule is surrounded by three OPG molecules, two glycerol molecules, one trehalose molecule and one urea molecule. Glycerol not only intermediates interactions between PMB1 and OPG, but also with 1-palmitoyl,2-cis-vaccenyl-phosphatidyl ethanolamine (PVPE) lipids, in this aggregate. PMB1 also interacts with PVPE *via* Dab6 and Dab10 - phosphate salt bridges. Additionally, the cluster was visited by two putrescine molecules – one remaining in the aggregate for 160 ns and the other for only 50 ns.

### PMB1 interaction with proteins

The number of interactions between PMB1 molecules and proteins were calculated based on intermolecular contacts (distances < 0.4 nm) during the course of the simulations, values concatenated over all trajectories for each protein are provided in Table S1. Interactions of PMB1 were observed with all of the different proteins in the systems. We consider the lipoprotein carriers, LolA and LolB first. In PMB_prot_ and PMB_crowd_ PMB1 molecules were found interacting both near to and at the entrance to the hydrophobic cavities of both proteins (Fig. 5 and 6). It is a useful reminder here that the normal function of LolA and LolB requires the lipid tails of cargo lipoproteins to bind in their hydrophobic cavities. A number of different PMB1 to LolA/B binding events were observed in our simulations. For example, in the PMB_prot_ system three molecules of PMB1 were observed to interact with the entrance to the cavity of LolA simultaneously (Fig. 5A), throughout one of the 500 ns simulations, with one partially inserted into the cavity. The LolA residues involved in theses interactions range from hydrophobic to charged: Trp49 (51%), Met51 (95%), Thr52 (69%), Gln53 (72%), Pro54 (46%), Asp55 (45%), Phe72 (32%) and Glu74 (25%), where parentheses indicate percentage of simulation time for which the interactions existed, reflecting the chemical diversity of PMB1. Interestingly, in PMB_prot_, one molecule of PMB1 bound to the entrance of the LolB cavity with its Lip1 residue inserted into the cavity (Fig. 6). Due to the extended conformation adopted by this PMB1, several LolB residues participated in long-lasting interactions (more than 60% of total simulation time) (Fig. 6B-C) with PMB1, including residues previously predicted as important for the binding of acyl chains^45^, namely Phe37, Val46, Met107 and Ile109. This indicates that PMB1 can bind in the LolB cavity in a manner that resembles normal the binding of acyl chains of lipoproteins to LolB.

**Figure 5:**
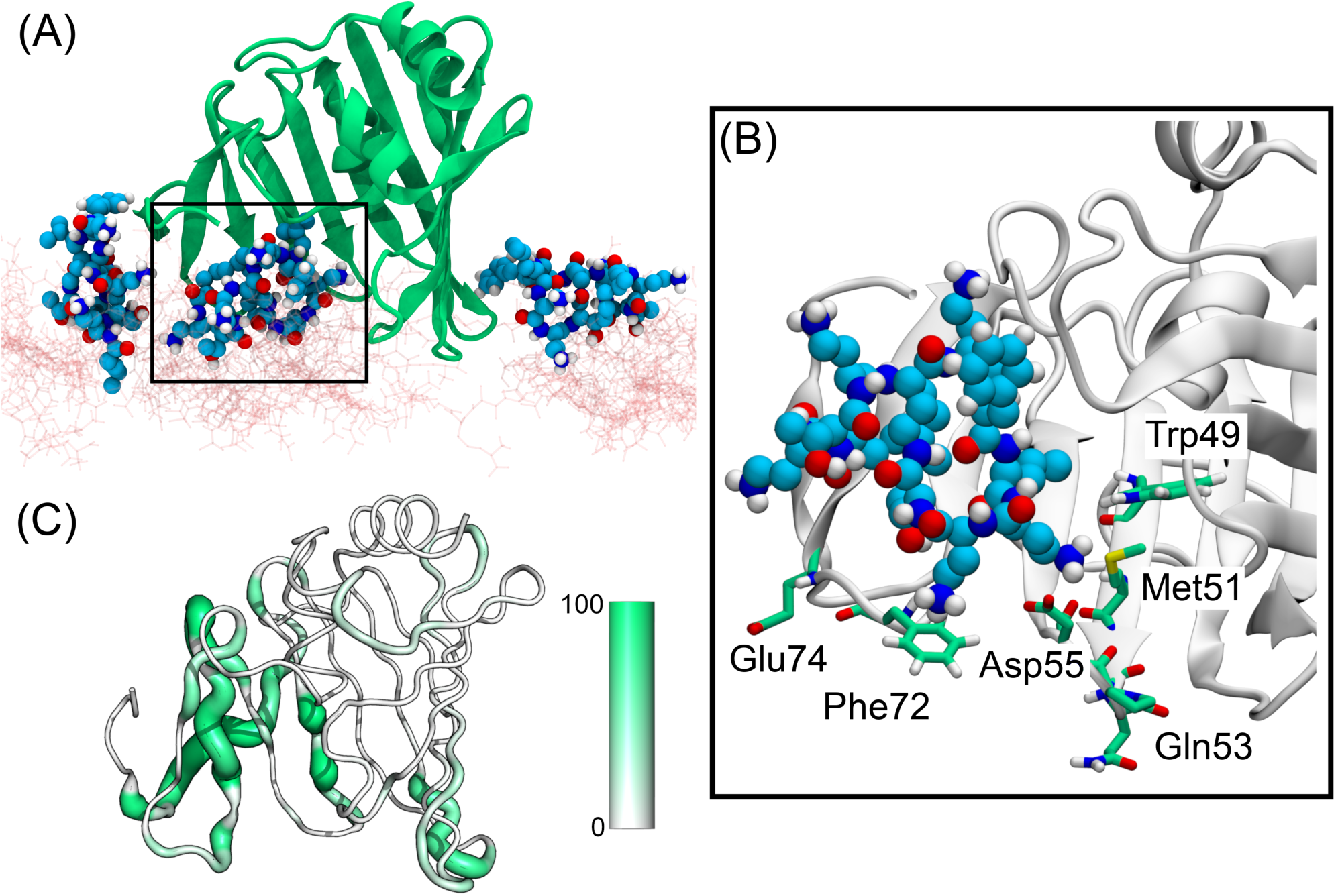
PMB1 binding modes to LolA. In PMB_prot_, three PMB1 bind to LolA at the same time (A) LolA = green, PMB1 = as previously, cell wall = pink sticks). One PMB1 is partially inside the hydrophobic cavity. (B) Zoomed in region where PMB1 is bound to the hydrophobic cavity of LolA, interacting mainly with residues Trp49, Met51, Gln53, Asp55, Phe72, and Glu74. (C) Sausage representation of LolA with respect to PMB1 interactions. Regions of the protein with higher percentage of time spent interacting with PMB1 are shown as enlarged tube, while regions with fewer interactions are shown as narrower tubes.

**Figure 6:**
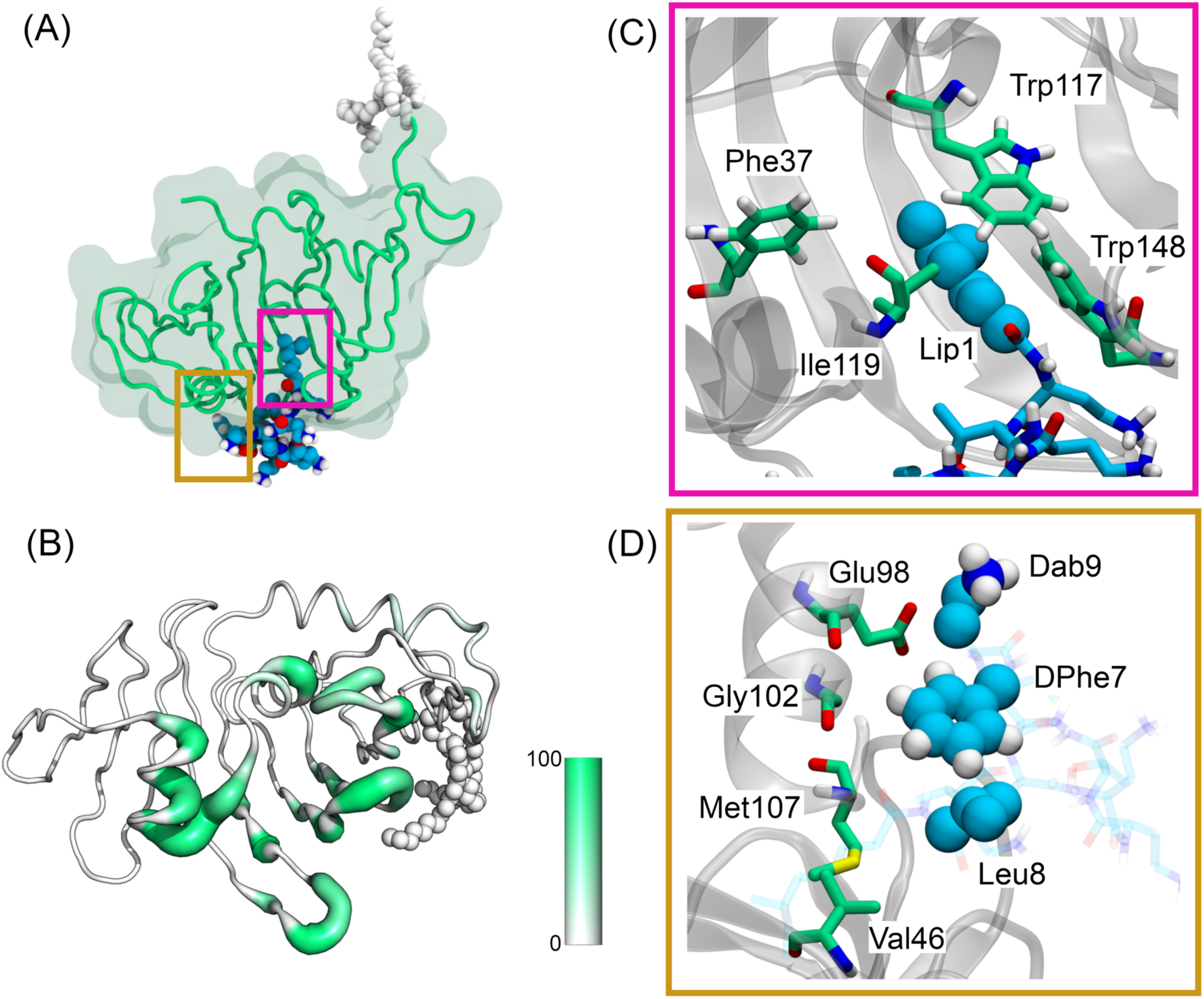
PMB1 binding mode to LolB. (A) PMB1 inserts Lip1 inside the hydrophobic cavity of LolB, reaching hydrophobic residues that usually interacts with acyl chains from lipoprotein ligands. (B) Sausage plot representation of the structure of LolB. Regions of the protein with higher percentage of time (from 0 to 1, coloured from white to green) spent interacting with PMB1 are shown as enlarged tube, while regions with less interactions are shown in thinner tubes. (C) Zoomed-in region showing binding at the hydrophobic cavity of LolB. In this area, Lip1 interacted with Phe37, Ile109, Trp117, and Trp148. (D) Zoomed-in region showing binding at the exterior part of LolB. In this region DPhe7, Leu8 and Dab9 interacted with residues Val46, Glu98, Gly102 and Met107.

Next we consider PMB1 - Pal interactions. In both the PMB1_crowd_ and PMB1_prot_ simulations, PMB1 was observed sandwiched in between the C-terminal domain (CTD) of Pal, the linker domain and the outer membrane. The interactions lasted for the entirety of the simulations. Pal residues involved in the interactions are provided in Table S1. Furthermore, upon PMB1 binding, the motion of the linker region of Pal become more restricted (Fig. S4), but the initial non-covalent interaction of Pal with peptidoglycan seems to be unaffected (Fig. S5). Despite that, this binding to the linker appears to limit the increase in number of contacts between Pal and the cell wall, when compared to systems that do not have a PMB1 attached to the linker (Fig. S5). More transient PMB1-Pal binding events also occurred in simulations of each system. In PMB_prot_, a PMB1 molecule entered the area in between the cell wall and the CTD of Pal (residues Ala109, Asp110, Arg112, Thr114, Tyr117 and Gly149), after initially being bound to a BLP for 370 ns (Fig. S6A). Other examples of PMB1 exchanging binding partners were also observed from Pal to LolA (Fig. S6B), from BLP to LolB (Fig. S6C), and from BLP to BLP (Fig. S6D). Thus, showing that within 500 ns PMB1 can move from interacting with one protein to another. In PMB_crowd_, after intermittently interacting with Leu176 and Lys185 in the Pal CTD for 160 ns, one PMB1 moved slightly away from Pal and associated with another PMB1 forming a dimer (Fig. S7). While the same region of Pal that was previously bound to the original PMB1 formed an interaction with a small cluster containing one molecule of OPG, one molecule of Glycerol and one molecule of putrescine. This small cluster also simultaneously interacted with the newly formed PMB1 dimer, which at this stage was not directly interacting with Pal.

PMB1 interactions with OmpA were mainly with the CTD. Each system had two or three PMB1 molecules binding to OmpA simultaneously. At least one molecule in each system was bound at the interface between the OmpA CTD and the cell wall, mediating the interaction between both structures. This binding region was located between the two main helices (composed of residues Glu212 to Asn226 and Ser253 to Lys267) from the CTD, with a prominent role of residues Gln214 (92%) and Tyr263 (87%). Interactions were hydrogen-bonding (Dab9 - Gln214) and hydrophobic (DPhe7 and Leu8 – Tyr263) in nature. Pal and OmpA have some structural similarities in their C-terminal domains (similarity of 35%), and both are known to bind to the cell wall^46-48^. Analysis of the contact data between PMB1-Pal and PMB1-OmpA revealed two long-lived interactions involving a specific helix from the CTD of each protein. This helix is composed of residues H_112_ANFLRSNPS_122_ in Pal and Y244SQLSNLDP252 in OmpA. Interestingly this region forms part of the dimerization interface of OmpA, thus would only be available for interaction when the protein is in its monomeric state^16,48^. Additionally, PMB1 was observed to bind to these regions while simultaneously interacting with adjacent motifs (Fig. S8) in both proteins: Q_63_MQQLQ_68_ in Pal (a short helix) and K_290_GIPADKIS_298_ (a loop connecting an α-helix to a β-strand). Interestingly only one PMB1 across all simulations was observed binding directly to the linker region of OmpA (in PMB_prot_), in PMB_crowd_, the linker area is largely occupied by osmolytes.

Comparison of data from PMB1 binding to BLP and Pal, revealed a short motif comprising the sequence S-S-E/D/N-X-Q/N (Fig. S9). Serine and acidic residues have particular propensity to interact with PMB1 (Fig. S10) due to the possibility of interacting *via* hydrogen bonding or salt bridges. Interactions between these residues are PMB1 lasted from 20 ns up to 400 ns. In simulations containing proteins, the four BLPs were the main target of binding events, with an average of 6 molecules of PMB1 binding to the four BLPs per system simulated. Regions of longest interactions (over 70% of simulation time) include residues S_11_SDVQTLNA_19_ (at the vicinity of the cell wall) and Asp34, Asp41, Ala42, Ala43, Arg48 (adjacent to the outer membrane). A number of the molecules interacting with BLP were also inserted in the interface with either the cell wall or the outer membrane in all of the simulations. Interestingly, in the PMB_crowd_ system most of the PMB1s that interacted with BLP were located in the solution region. The interfacial regions favoured by PMB1 in other systems (PMB_dil_ and PMB_prot_) were occupied by osmolytes in PMB_crowd_. Thus, the presence of osmolytes seems to force PMB1 into the solution to some extent, by occupying interfacial binding regions.

### PMB1 interactions with the cell wall

A number of PMB1 molecules (at least 14) reached the peptidoglycan layer area very rapidly (within the first 10 ns) in all simulations (Fig. S1 and S11). The negative charges presented by D-Glu and meso-diaminopimelate (m-DAP) residues from the peptide portion of the cell wall interact with PMB1. Other interactions are also present, such as PMB1 forming hydrogen bonds to the hydroxyl groups attached to the pyranosidic rings of the peptidoglycan glycan strands (Fig. S12). None of the PMB1 molecules were observed to go through the pores of the cell wall and dissociate from it during the total time of 3 μs of all simulated systems. Most osmolytes also did not cross through the pores easily: only putrescine, trehalose and urea were able to cross multiple times (Fig. S13). Generally, two modes of association between PMB1 and the cell wall were observed. In one mode, PMB1 inserts itself between glycan strands, as seen in one example from PMB_dil_ system (Fig. 7A). In some cases, it acted similarly to a peptide linkage, as it was able to form salt bridge interactions with both glycan strands simultaneously for more than 200 ns (Fig. 7C), decreasing the local distance between the strands from ∼2.7 nm to ∼1.9 nm. The other observed mode of association is PMB1 attaching to the surface of the cell wall, not inserted between the strands, but rather located around peptide linkages (Fig. 7B). A common aspect from both binding modes is that the Dab residues from PMB1 attract the loose peptide portions (not connected to > 1 glycan strand), forming salt bridges. The insertion mode of interaction involved the formation of multiple long-lived (∼ 200 ns) salt bridges with the Dab - m-DAP and Dab - D-Glu, in contrast, in the surface binding mode the salt bridges had a lifetime of ∼ 100 ns. (Fig. 7D).

**Figure 7:**
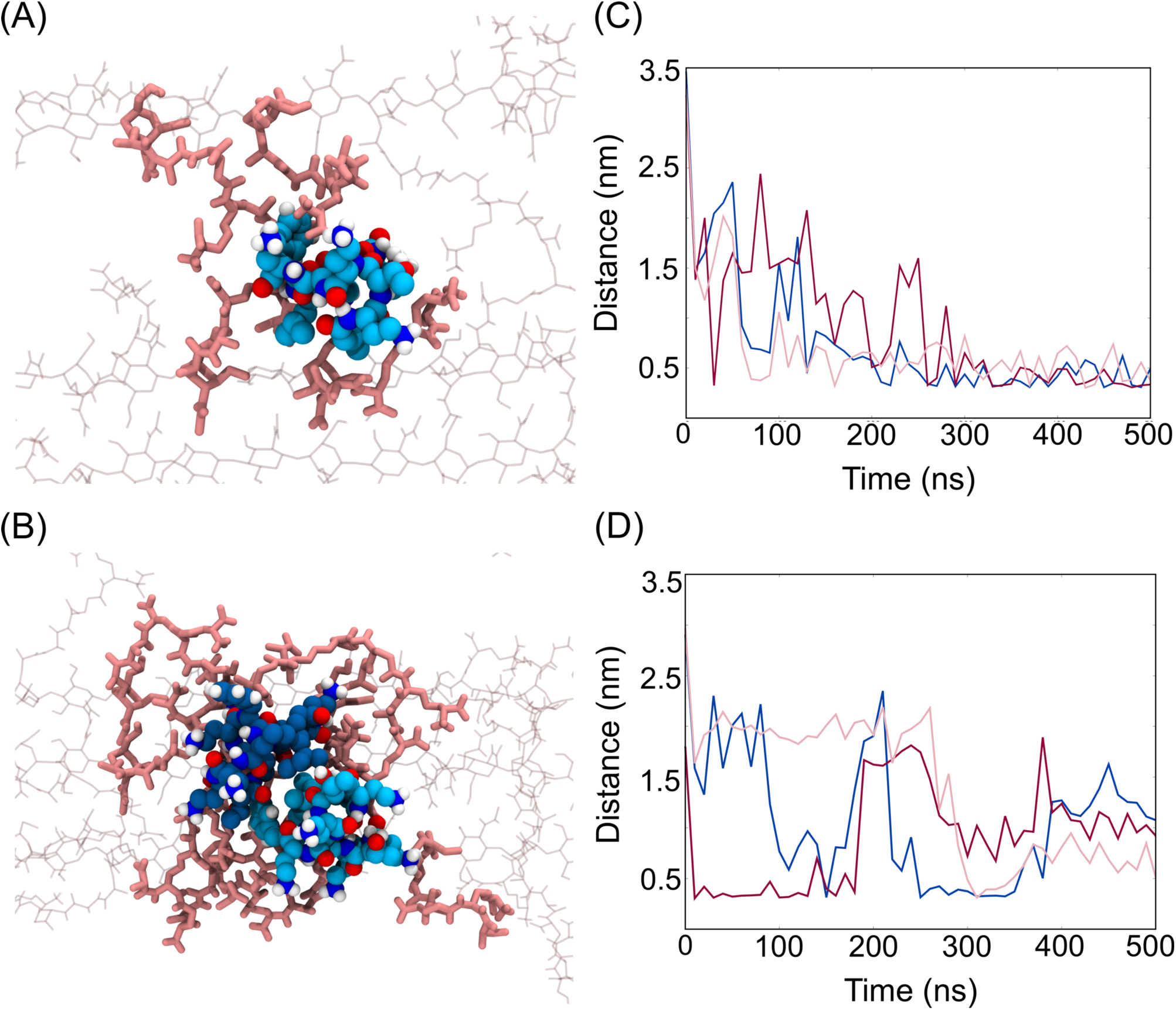
PMB1 binding modes to the cell wall extracted from PMB_dil_ and PMB_prot_. The inserted binding mode is depicted on (A), showing PMB1 (blue carbon spheres) attached inside of the pores of the cell wall (pink sticks) and interacting with many negatively charged residues. The surface binding mode is depicted in (B), showing a PMB1 dimer (two shades of blue) bound to the surface of three glycan strands. (C) and (D) shows examples of distances of salt bridges interactions between different Dab residues (PMB1) and m-DAP and D-Glu (cell wall). In the inserted binding mode, interactions seem to last longer overall than in the surface binding mode.

In the PMB_prot_ system, a few of the PMB1 molecules mediated protein binding to the cell wall (Fig. 8). For example, PMB1 bound to the surface of LolA, interacting with residues Lys23, Asp26 and Glu34, while at the same time interacting with m-DAP and D-Glu from the cell wall. An observation from both simulations of the PMB_prot_ system, was an extension of the OmpA linker region that appeared to be induced by PMB1 which is bound to the CTD region. By around 420 ns, there is an increase in the number of residues from OmpA contacting the cell wall, increasing from 5 to 10 (Fig. S14). This appeared to occur spontaneously, but subsequently the PMB1 Dab residues form salt bridges with three charged groups (two D-Glu and one m-DAP) of the loose peptide ends of the cell wall (Fig. 8 and S14). These interactions of PMB1 which is bound to both OmpA and the cell wall, seemed to have acted as a driving force to push the OmpA CTD further towards the cell wall, stabilizing and increasing the OmpA-peptidoglycan binding interface from 10 residues to almost 20. Similarly, in the PMB_crowd_ system, some associations between the PMB1, proteins (OmpA, LolA and Pal) and the cell wall were also observed. For example, initially, a PMB1 is associated with the OmpA CTD mediated by two trehalose molecules and one glycerol molecule (Fig. 8). At the same time, the same PMB1, one of the trehalose, glycerol and residue Asn249 of OmpA were in the vicinity of the cell wall (within 0.6 nm), displaying a few hydrogen bonding interactions. By the end of the simulation, this cluster had dissociated. Another example is the PMB1-Pal association mentioned in the previous section with the presence of a small cluster of osmolytes (Fig. 8). Residue Asp179 of Pal interacted with OPG, glycerol, and putrescine molecules while they were bound to PMB1 and/or the cell wall, the interaction partners in this cluster changed frequently over time, indicating the non-specificity of these intermolecular associations.

**Figure 8:**
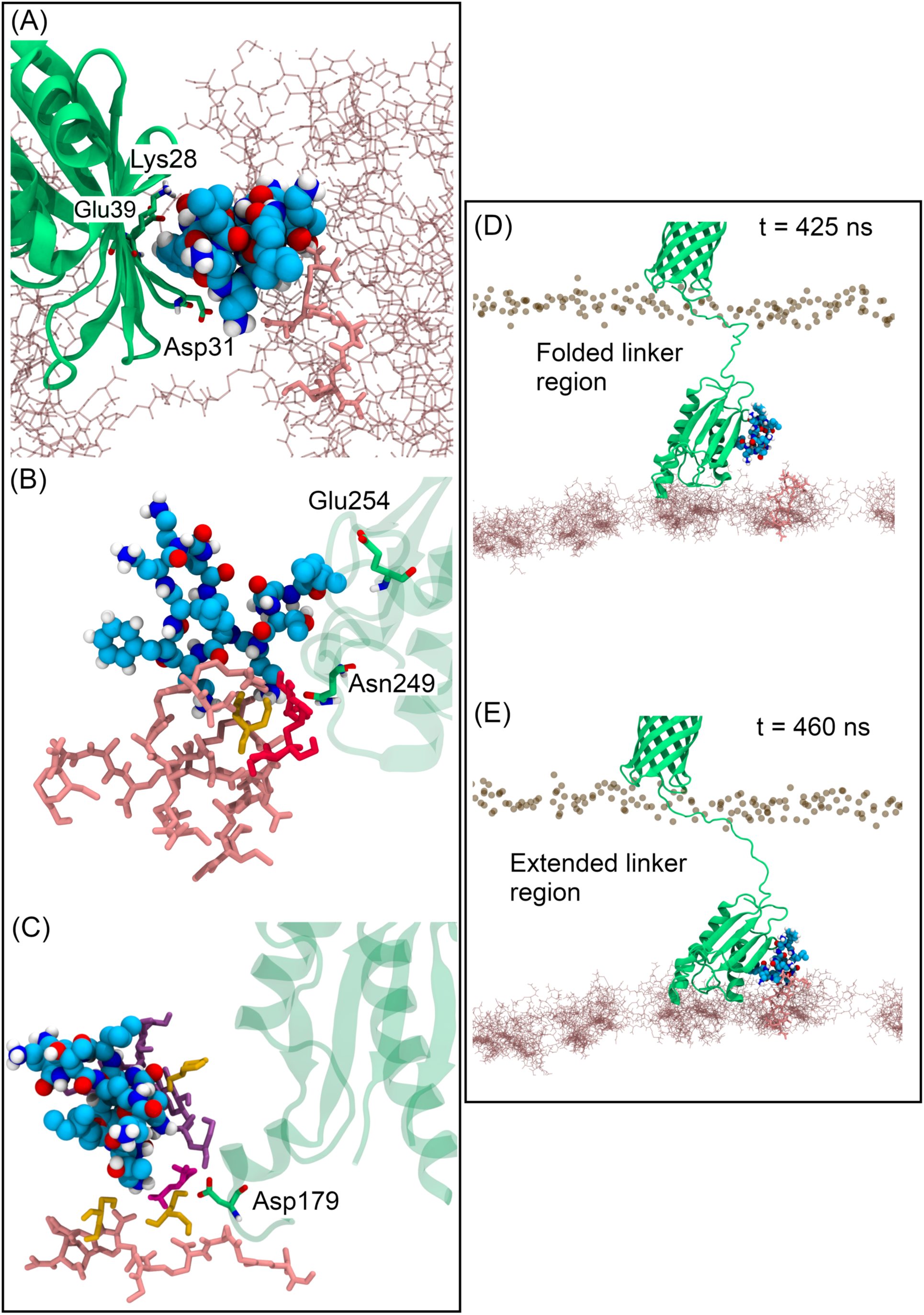
PMB1 mediating interactions between LolA, Pal, OmpA (green) and the cell wall (pink sticks). Panel (A) shows highlighted residues Asp26, Lys23 and Glu34 (green) from LolA interacting with Dab (Asp26) and DPhe7 (Lys23 and Glu34,) from PMB1 (colored in blue spheres), while another Dab from PMB1 interacts with a negative charged residue from the cell wall (pink licorice sticks). In Panel (B), PMB1 and osmolytes (glycerol – orange; trehalose – red) are shown mediating interactions for OmpA-cell wall. In (C), a bigger cluster containing OPG, putrescine, glycerol and urea is shown mediating interactions with Pal and the cell wall. Panels (D) and (E) depict two different states of OmpA and PMB1, before (D) and after (E) enhancing the interaction with the cell wall. In (B) and (C), only cell wall residues that are in contact with PMB1 and osmolytes are shown to improve clarity.

## Discussion

Currently there is a great need to find novel therapeutic agents to address the problem of antimicrobial resistance to antibiotics^49,50^. In order to do so in a rational manner requires a thorough understanding of the environment faced by antibiotics such as PMB1 as they negotiate the bacterial cell envelope. In this work, we have simulated an atomistic model of the periplasmic space to study the fate of PMB1 in this region once it has already crossed the OM. Our results predict that PMB1, and likely other drugs relying on diffusion alone to cross the periplasm, face a complex path, full of molecular obstacles which hinder their movement through the periplasm. The presence of structures from subcellular compartments in our model systems had a major effect on the diffusion coefficients of PMB1 molecules. This was observed by the 100-fold reduction in values in the periplasm models compared to PMB1 just in solution. Similarly, diffusion of native proteins was also affected by increased crowding. Previous studies of diffusion rates in the cytoplasm and the periplasm showed that the GFP proteins have a slightly slower diffusion rate in the periplasm^38^ when compared to the cytoplasm and there have been some discussions in the literature regarding the nature of the periplasm being a gel-like environment^51^ or a fluid environment^38^. At the level of crowding we have in our simulations the system is clearly still fluid. Crowding up to 30% of volume has been reported previously to have modest effects on water properties^34^, showing alterations in the self-diffusion coefficient of water that are in-line with our simulations (∼20% Volume with ∼2.0 × 10^−5^ cm^2^/s).

PMB1 is an amphipathic molecule with considerable conformational flexibility. In just water and ions, we observed PMB1 tetramers forming micelles with sizes comparable to experimental data^30^ for micelle formation in solution of colistin (polymyxin E). In addition, as predicted for colistin and its analogue^39^, our micelle formation followed a “closed association” model, in which the number of monomers is discrete, limiting micelles to a certain size (pentamers with 2.6 nm). Thus, our model of PMB1 in water provided a ‘baseline’ reference system which gave aggregate sizes and dynamical behavior that reproduced experimental observables for similar molecules.

We observed a wide range of associations of PMB1 with other molecules. One particularly interesting phenomenon observed here is PMB1 insertion into the hydrophobic cavities of the lipoprotein carriers LolA and LolB. These cavities have previously been shown to be non-specific binders of hydrophobic molecules^52-54^. Our results suggest it is possible that some PMB1s may be carried through the cell wall by hijacking the lipoprotein carrying functionalities of LolA/LolB. Given we observe PMB1 adhering to the cell wall this ‘hitchhiking’ mechanism would be advantageous in providing an easier route through the cell wall. To our knowledge this spontaneous phenomenon is described here for the first time. It is important to consider here other potential consequences of PMB1 binding to the lipoprotein carriers. LolA and LolB play important roles in avoiding toxicity due to accumulation of BLP in the inner membrane^55,56^. Binding of PMB1 into their hydrophobic cavities may serve to inhibit their natural functions and lead to mislocalization of lipoproteins in the inner membrane, in a similar manner to small hydrophobic inhibitors such as MAC13243^52-54^. Interestingly, LolA transcription is triggered with increasing concentrations of PMB, a mechanism connected to the activation of the stress regulator Rcs phosphorelay system^57^ which provides indirect evidence to support our hypothesis. Osmolyte - PMB1 interactions varied depending upon the chemistry of the osmolyte. We observed fast formation of small clusters of molecules, with PMB1 usually binding to, and often becoming partially coated with the polar OPG and glycerol molecules. The OPG concentration becomes slowly diluted when bacteria are moved to concentrated media^58^. Given the high propensity for these molecules to bind to PMB1 in our simulations, we suggest that the changes in the OPG concentration may also impact on dynamics of PMB1 in the periplasm. As discussed in previous simulation studies^34,35,37^, molecular crowders can promote a range of effects, including excluded volume effects and replacing interactions. In our simulations, osmolytes mediate interactions between other molecules and also replace some interactions. For example in the absence of osmolytes, there is a greater propensity for PMB1-BLP interactions to occur at the BLP/cell wall and BLP/OM interfaces, however these regions are occupied by osmolytes in the most crowded system, and consequently PMB1 interactions with BLP are largely with the region of the protein in ‘bulk’ solution in the periplasm. This suggests that the non-specific binding of osmolytes to cell envelope components may have local consequences for available binding modes for antibiotics.

Two main peptidoglycan binding modes were observed; one in which PMB1 inserts in between glycan strands acting as a pseudo cross-link and one in which it is surface-bound close to the peptide linkages. Atomic Force Microscopy (AFM)^59^ studies have suggested that the cell wall interactions formed by colistin may be responsible for rigidifying the cell envelope.

A limitation of our results involves the correction of the diffusion coefficients for finite-size effects. Given the complexity of our models, with the presence of the outer membrane and the cell wall, that our simulation boxes are non-cubic, and that the correction becomes smaller with increasing box size, we opted for not applying it in these D_t_ values. Our results for proteins and water diffusion correlate well with previous atomistic crowding models^34-37^. Experimental studies^38,60^ reported D_t_ rates for GFP that are in a similar range to ours. We note here that experimental value for the OmpA N-terminal domain (transmembrane) alone, D_exp_ = 4.9 ± 0.09 × 10^−7^ cm^2^/s is faster than the value obtained for the complete protein from our simulations (D_t_ = 3.1 ± 0.6 × 10^−8^ cm^2^/s in PMB_prot_ and D_t_ = 1.8 ± 0.5 × 10^−8^ cm^2^/s in PMB_crowd_). The C-terminal domain of OmpA is bound to the peptidoglycan which is highly likely to be the cause of the slower diffusion in the simulations.

It is also worth mentioning that the SPC water model (which works well with the GROMOS54a7 force field^61^) overestimates experimental values of self-diffusion for water molecules^62^, so D_t_ values for solutes possibly are affected by this. Finally, crowding systems simulations are very complex and could lead to intense aggregation when using standard additive force fields^63,64^, with several methods being proposed to solve this^65,66^. From our perspective, GROMOS54a7 was a reasonable choice, since it is one of the force fields that shows a lesser preference for the aggregated state^70^, while also having validated parameters for the complex mixture of lipids that compose the bacterial outer membrane and the bacterial cell wall. Our results where comparable with other experimental and simulation studies are in-line with those, providing further confidence in the predictions from the complex simulations which go beyond previous studies in terms of the resolution and complexity of the simulations studied.

## Conclusions

In conclusion, atomistic molecular dynamics simulations of the antibiotic PMB1 in a number of models of the periplasm of Gram-negative bacteria with differing levels of crowding have revealed slower diffusion of the antibiotic as the periplasm becomes more crowded. PMB1 forms complexes with osmolytes, the cell wall and native cell envelope proteins which can be short-lived or long-lived. PMB1 is rarely uncomplexed within the periplasm, therefore its functional groups are occupied in interaction with other species more often than not. We feel this is an important factor to consider in future development of antibiotics (and may be extended to drugs that target other organisms too). The *in vivo* environment is not a chemistry experiment in which ones controls the type and number of molecules involved, and the complexity of the former may impact upon foreign molecules such as drugs in many unexpected ways. The simulations described here show that incorporation of the chemical details of the local environment can predict likely interactions with other species and highlight potential mechanistic pathways that may have been originally unintended (such as the ability of PMB1 to bind to LolA and LolB).

## Methods

### System Preparation

We constructed the template model based on previously published works from our group^17,48,67^. This was composed by an asymmetric outer membrane (OM) of an identical composition as seen in previous works^17,67-69^, a one-layer peptidoglycan cell wall (PGN) formed by 12 glycan strands of 17 repeating NAG-NAM-peptide units, four Braun Lipoproteins (BLP) covalently attached to PGN and inserted in the membrane by tripalmitoyl-S-glyceryl-cisteine residues^67,69^. Models for BLP(PDB: 1EQ7)^70^, LolA (PDB: 1IWL) and LolB (PDB: 1WLM)^45^, OmpA^16^, Pal (PDB: 2W8B)^71^ were taken from previous works from the group^72,48,54,67^. Pal and OmpA proteins were also included in PMB_prot_ and PMB_crowd_ systems. Crystallographic structures of LolA and LolB GROMOS 54A7 force field with the GROMOS 53A6OXY ether parameters^73^ were used for the construction of the tripalmitoyl-S-glyceryl-cysteines. After setting up this initial template system, we added the other components: Pal bound and unbound to the cell wall, PMB1 molecules, and ions concentrations. PMB1 GROMOS 54a7 parameters were obtained by using the Automated Topology Builder (ATB) server^74^. For OmpA insertions into the OM, we employed the *gmx membed* tool^75^, similarly to a previous report^67^.

For the construction of the PMB_crowd_ system, parameters for the molecular crowders were obtained using the ATB server, with the exception of OPG and trehalose, in which the GROMOS 56a6 (CARBO)^76^ parameters were employed, which are compatible with GROMOS 54a7. We adapted the “droplet methodology” from Bortot *et al*^37^ to deal with the insertion of osmolytes, by adding each osmolyte with a water shell obtained from 100 ns molecular dynamics simulations.

### Atomistic Molecular Dynamics Simulations

We performed molecular dynamics simulations employing the GROMACS simulation suite (version 2018.3)^77^ along with GROMOS54a7 force field^61^ and SPC water model^33^. We divided the simulations in two parts: equilibration simulations in NVT and NPT ensembles with position restraints in proteins, cell wall, and PMB1, which lasted for 1 and 100 ns, respectively; and production simulations in NPT ensemble, which ran for 500 ns. Simulations were performed at 310 K, which was maintained by employing the velocity rescale thermostat^78^ with a coupling constant of τ = 0.1. Pressure was maintained semi-isotropically at 1 atm by employing the Parrinello-Rahman barostat^79^ with a time constant of 2 ps. The particle mesh Ewald method treated long-range electrostatics^80^. LINCS algorithm^81,82^ constrained the covalent bonds, which allowed an integration step of 2 fs. Both long-range electrostatics and van der Waals cutoffs were set to 1.4 nm. To neutralize charges, we added the correct number of counterions together with an extra salt concentration of 0.2 M of sodium chloride ions for all simulations. For the replicates, starting positions of the proteins and PMB1 molecules were changed, along with re-solvation of the system, equilibration and production phases. In addition, we modified the starting velocities to ensure the difference between runs and improve conformational sampling. For molecular manipulation, visualization, and analysis, we employed the VMD software^83^.

### Analysis

Translational diffusion coefficients, D_t_, were obtained by using the *gmx msd* analysis tool from the GROMACS tool set to calculate the mean square displacements (MSD). MSD plots (Figure S2) were calculated for time delays The linear fit (where the slope was obtained) was performed in different time regimes (1-10 ns; 50-100 ns) for PMB1 and water (1-10 ns; 10-100 ns) aiming to capture the slowdown in D_t_ due to crowding effects, while for proteins, the linear fit was performed in the range of 5-15 ns. Error estimates were obtained by averaging over all and by averaging over replicate simulations of each system. For the XYZ motion analysis, the *gmx trajectory* tool was employed to obtain the coordinates of the center of mass of each PMB1 in the X, Y and Z axis. Radial distribution function values (RDF) were obtained using the *gmx rdf* tool, while *gmx sasa* was used for SASA calculations. In-house scripts were employed for the intermolecular contact analysis.

## Supporting information

all support info

## ASSOCIATED CONTENT

### Supporting Information

The Supporting Information is available free of charge on the ACS Publications website. Further analysis of the data including Figures S1-S14 and Table S1 (PDF file).

### Author Contributions

The manuscript was written through contributions of all authors. SK conceived the project, CP conducted the simulations, IPSS and AB helpeed with simulation setup and analysis, CP and SK also performed analysis, CP and SK wrote the paper. All authors have given approval to the final version of the manuscript.

## Acknowledgements

The authors acknowledge the use of the IRIDIS High Performance Computing Facility, and associated support services at the University of Southampton, in the completion of this work. This project made use of time on ARCHER granted via the UK High-End Computing Consortium for Biomolecular Simulation, HECBioSim (http://hecbiosim.ac.uk), supported by EPSRC through grant number EP/R029407/1, which also supports CP.

## For Table of Contents Only

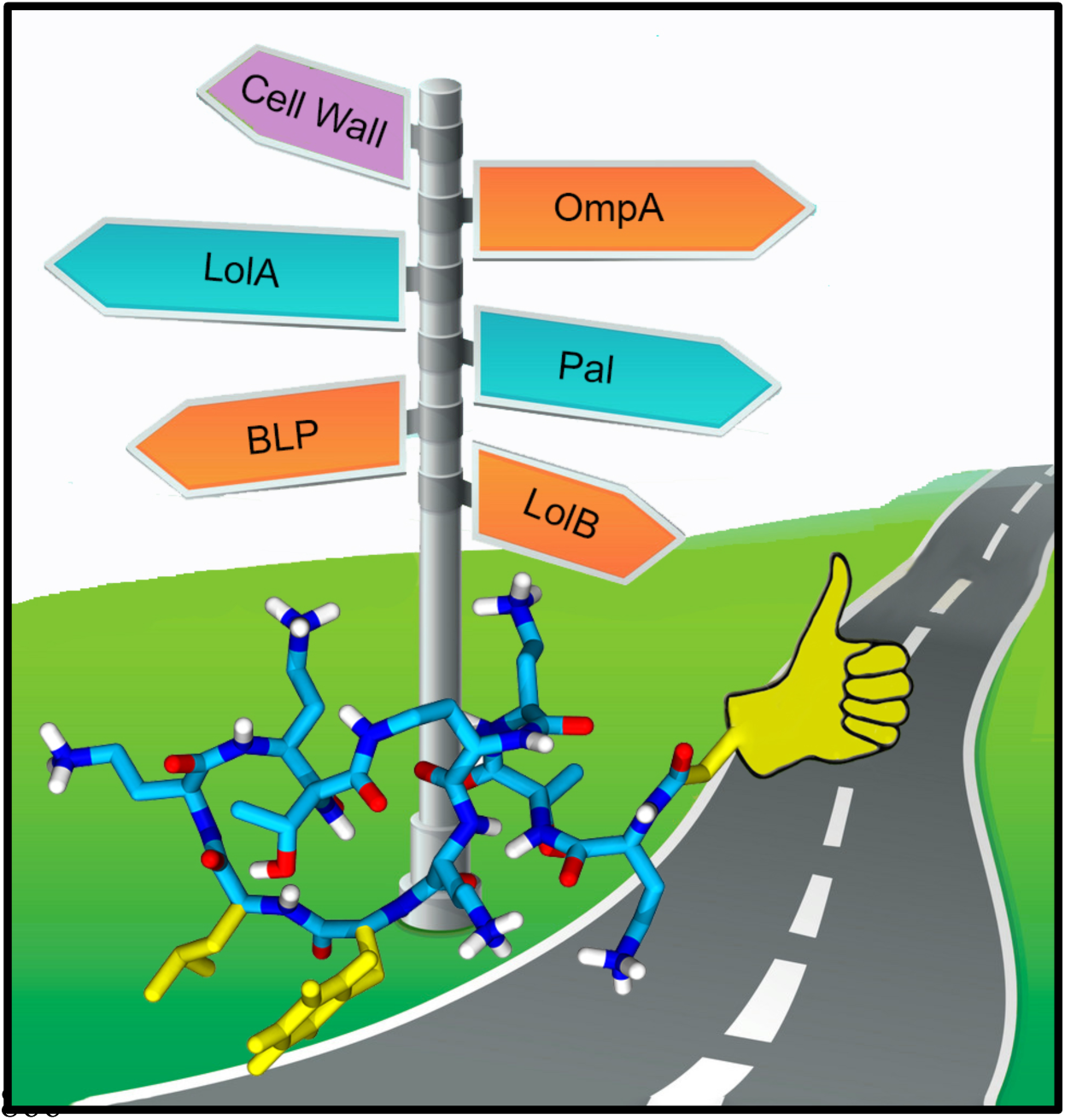

