## Supplementary material for "The Hitchhiker’s Guide to the Periplasm: Unexpected Molecular Interactions of Antibiotics Revealed by Considering Crowding Effects in *E. coli*": all support info

### Supporting Information for "The Hitch-hikers Guide to the Periplasm: Unexpected Molecular Interactions of Antibiotics Revealed by Considering Crowding Effects in *E. coli*"

Conrado Pedebos[1], Iain P. S. Smith[1], Alister Boags[1,2], Syma Khalid[1*]

[1]School of Chemistry, University of Southampton, Highfield Campus, Southampton SO17 1BJ, UK
[2]Bioinformatics Institute (A*STAR), 30 Biopolis Street, #07-01 Matrix, Singapore 138671, Singapore

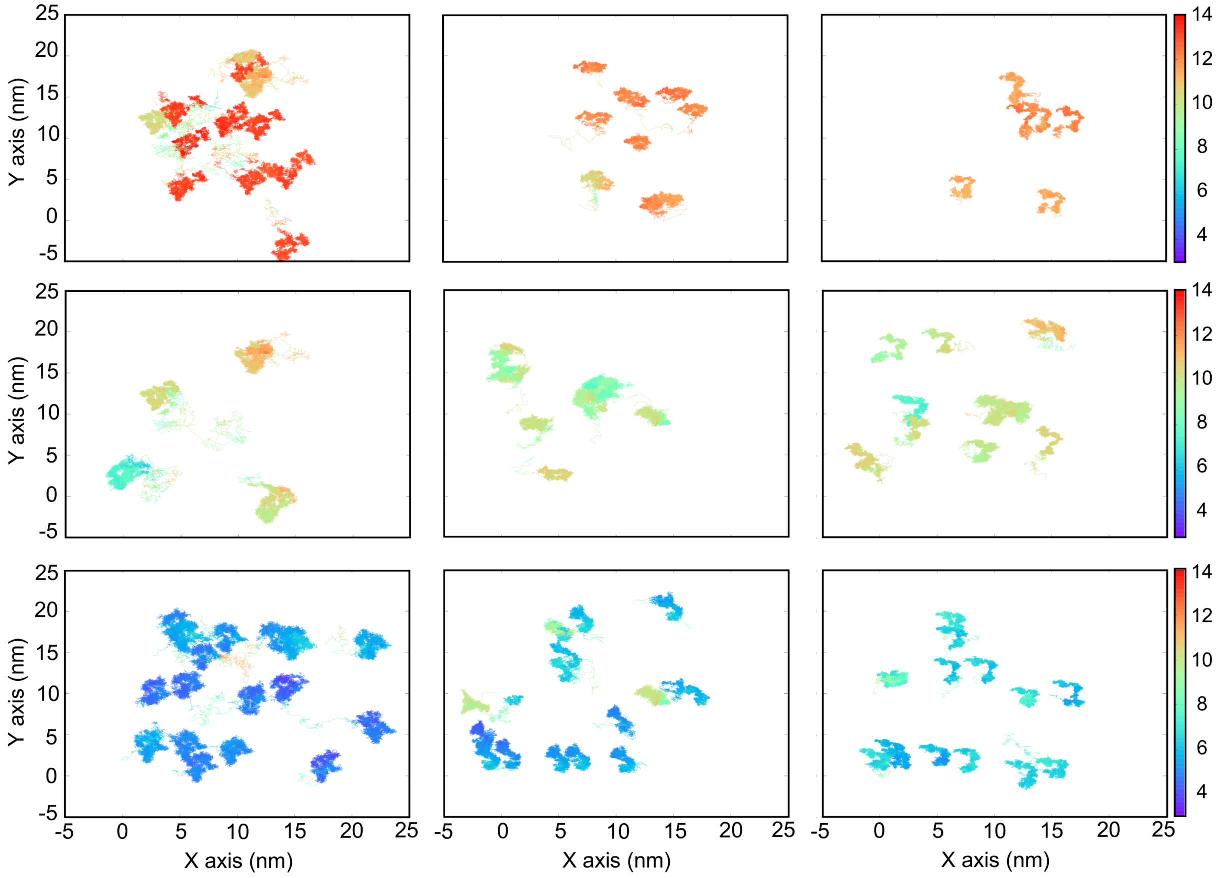


**Figure S1. PMB1 molecules motion in different areas of the system. Panels (A-C) show PMB1 motion in $PMB_{dil}$, (D-F) show PMB1 motion in $PMB_{prot}$, and (G-I) show PMB1 motion in $PMB_{crowd}$. Top row (A, D, G) depicts motion the outer membrane region. Middle row (B, E, H) depicts motion the solution region. Bottom row (C, F, I) depicts motion in the cell wall area. Z-axis range is described by the color bar.**

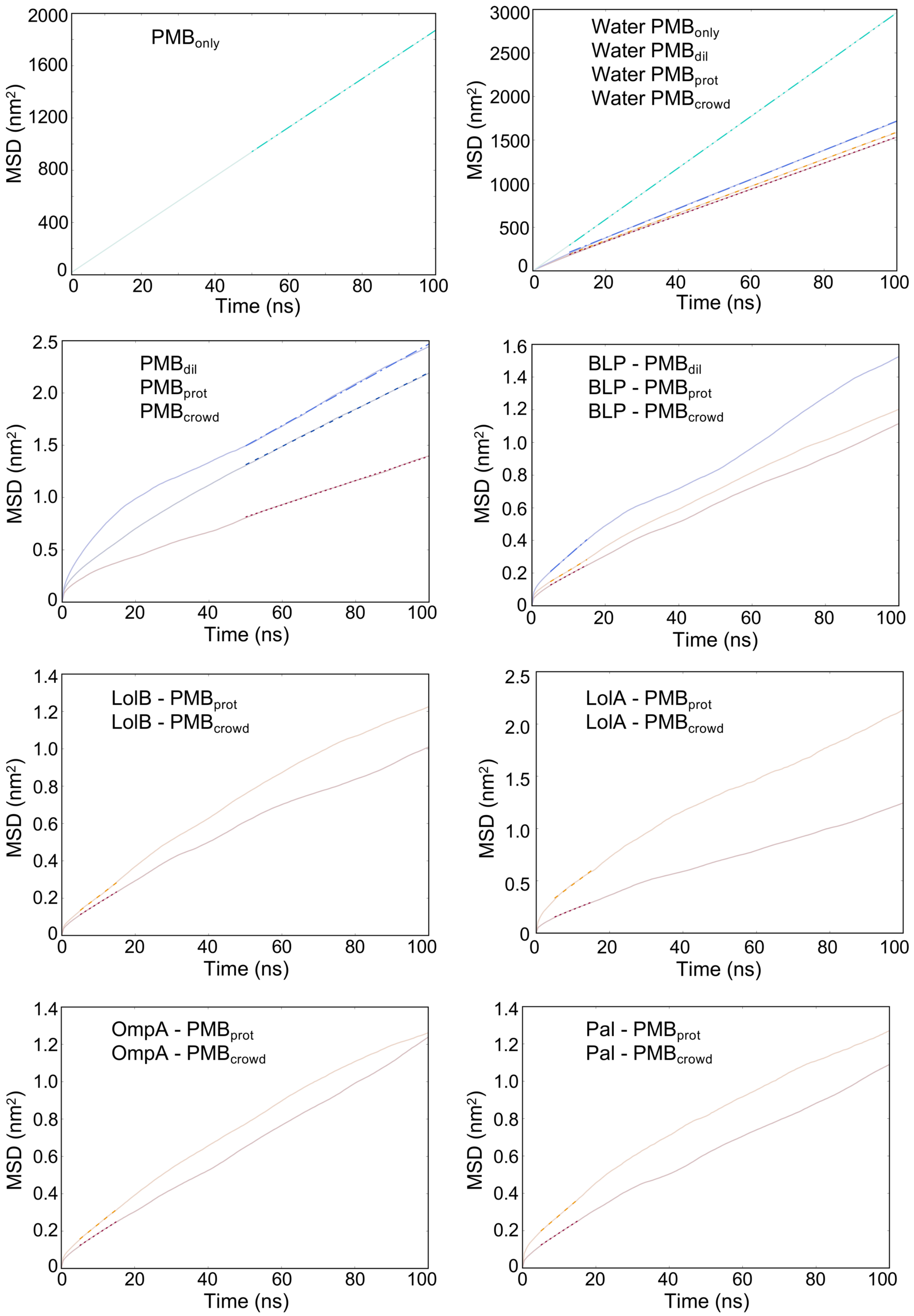


**Figure S2. Average MSD plots for the molecules of interest from this study. Each colored curve indicates a different system. Linear fitting is indicated by dashed**

**lines superimposed in the curves. Long-time regime fitted corresponded to 10-100ns for water; 50-100ns for PMB1; 5-15ns for proteins.**

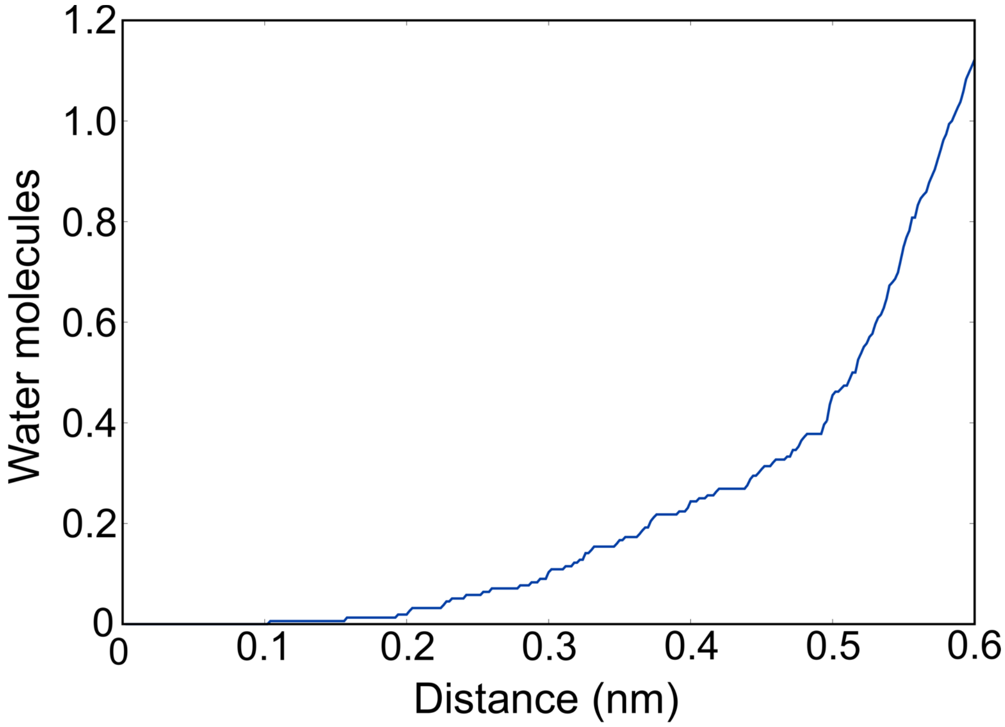


**Figure S3. Average number of water molecules within a distance cut-off of 0.6 nm from the center of mass of a tetrameric micelle, extracted from the radial distribution function data (cumulative number RDF).**

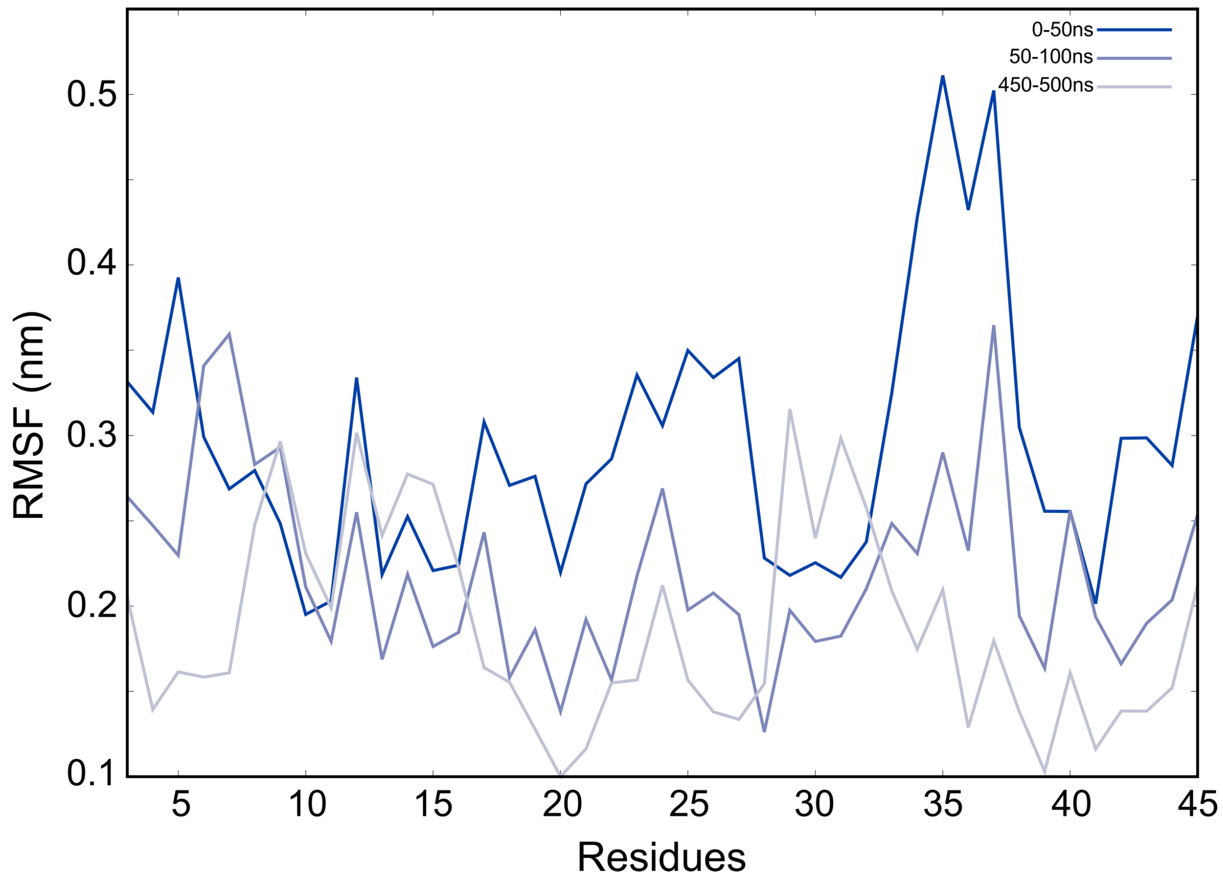


**Figure S4. Root mean square fluctuations (RMSF) of the linker region of the Pal protein in $PMB_{prot}$. Values are calculated for different timeframes, depicting the beginning and the end of the simulation and shown in the legend.**

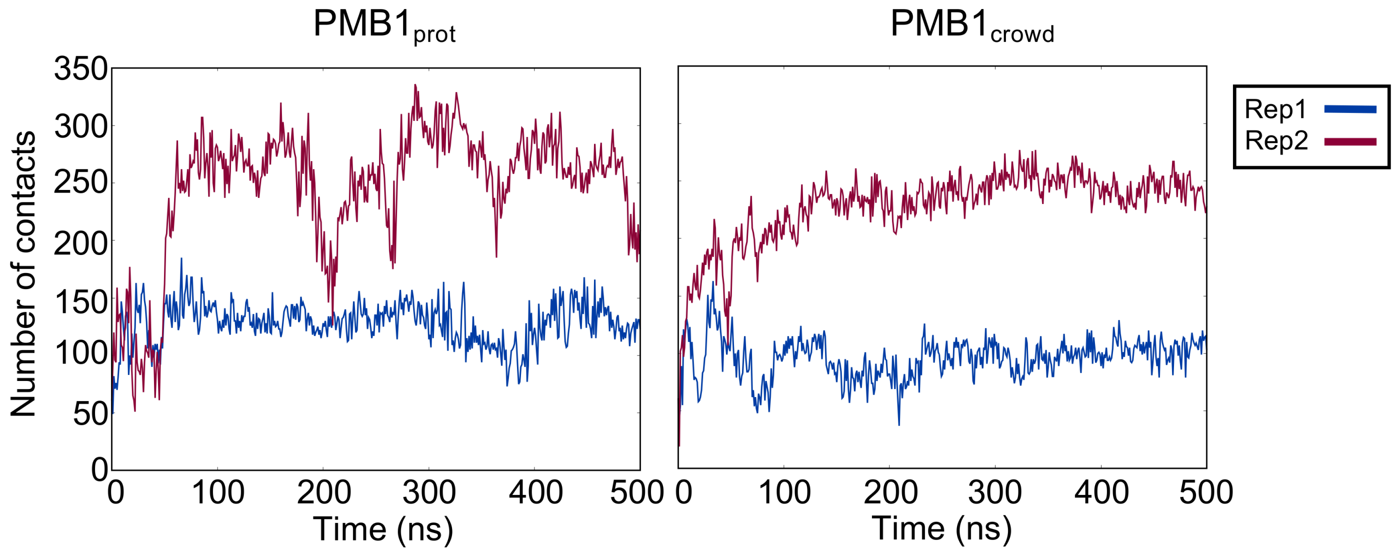


**Figure S5. Number of atomic contacts between Pal and the cell wall, using a cut-off of 0.4 nm. In both Rep 1 (blue curves) there is one PMB1 molecule attached to the linker region of Pal, while both Rep 2 (maroon curves) do not have any PMB1 bound to their linker regions.**

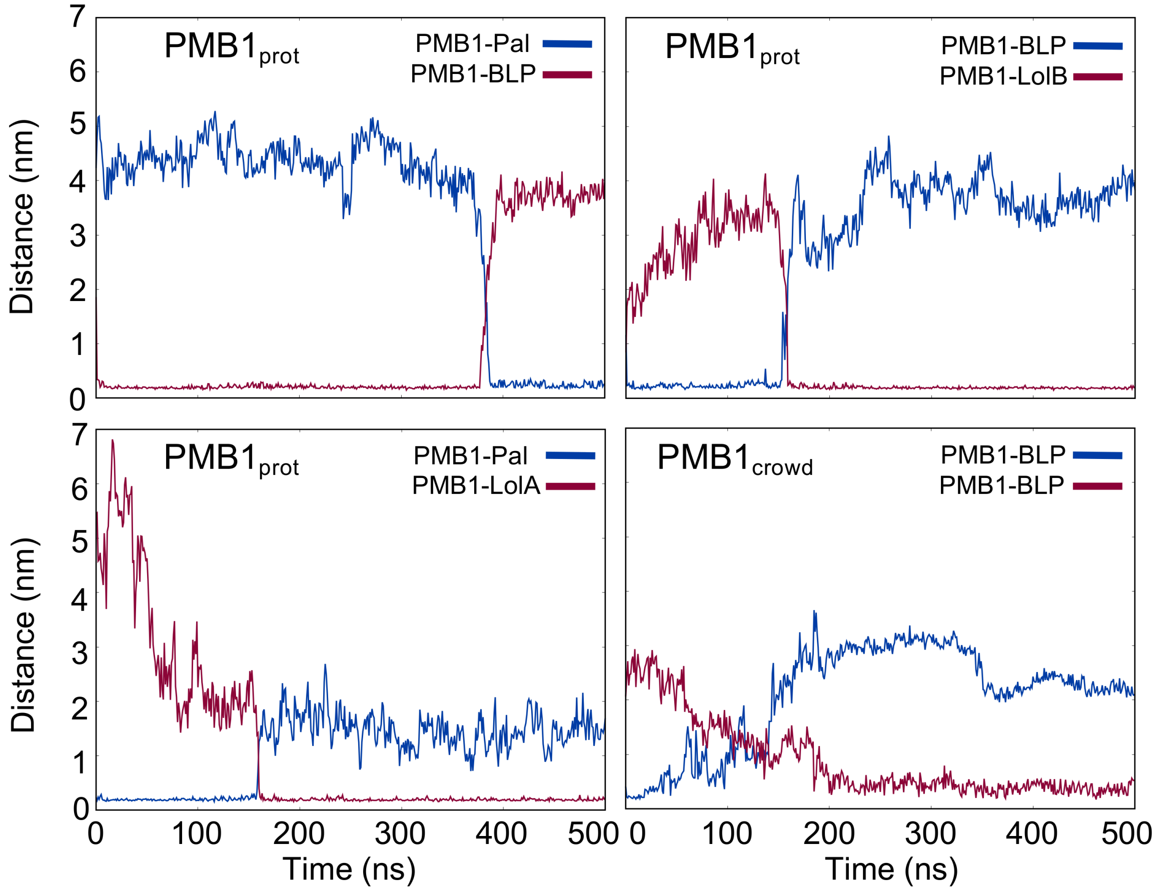


**Figure S6. Distance plots between PMB1 and proteins, displaying the transient binding of PMB1 molecules in the studied systems.**

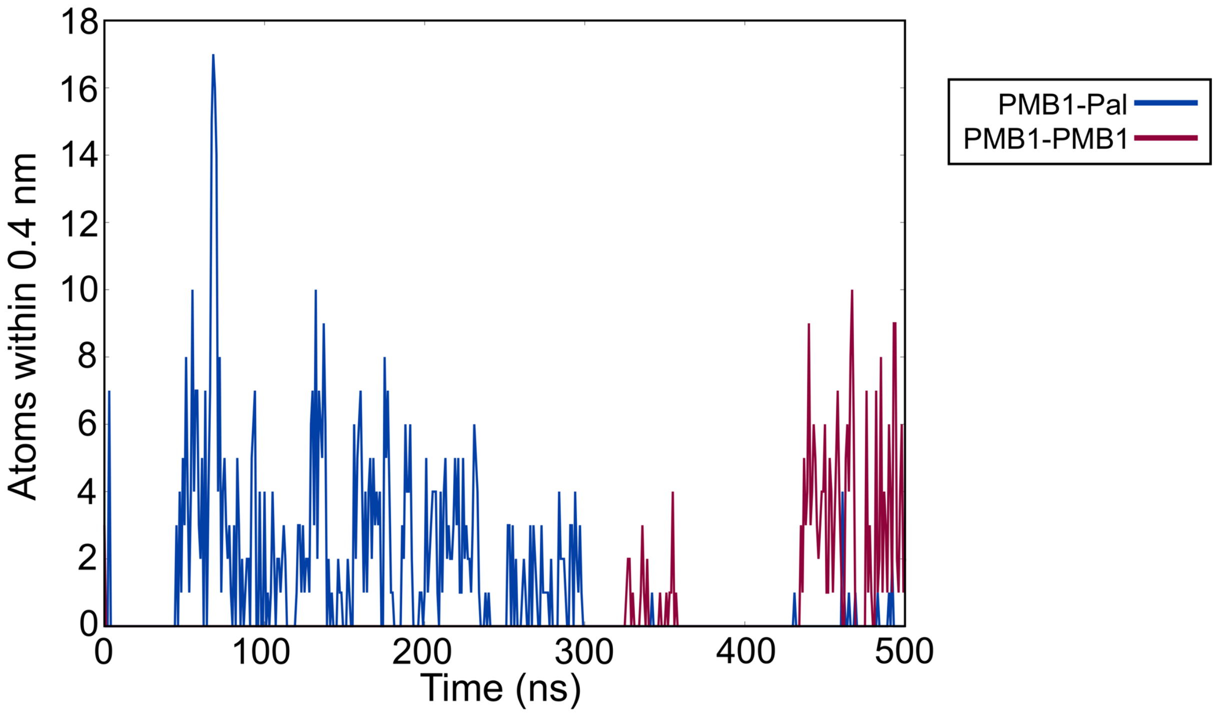


**Figure S7. Number of atoms within a distance cut-off of 0.4 nm, for PMB1-Pal (blue curve) and PMB1-PMB1 (maroon curve), showing the change in binding partners.**

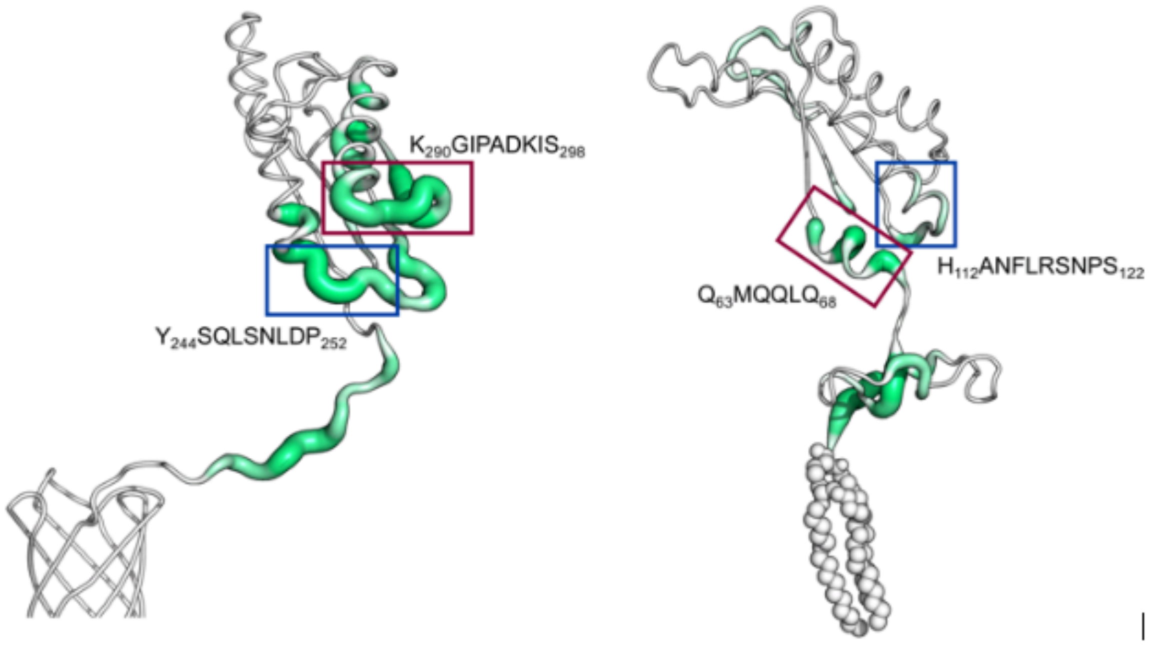


**Figure S8. Sausage plots of OmpA and Pal in PMB$_{prot}$. Regions of the protein with higher percentage of time spent interacting with PMB1 are shown as enlarged tube, while regions with fewer interactions are shown as narrower tubes. Similarly arranged regions in Pal and OmpA are highlighted with blue and maroon rectangles.**

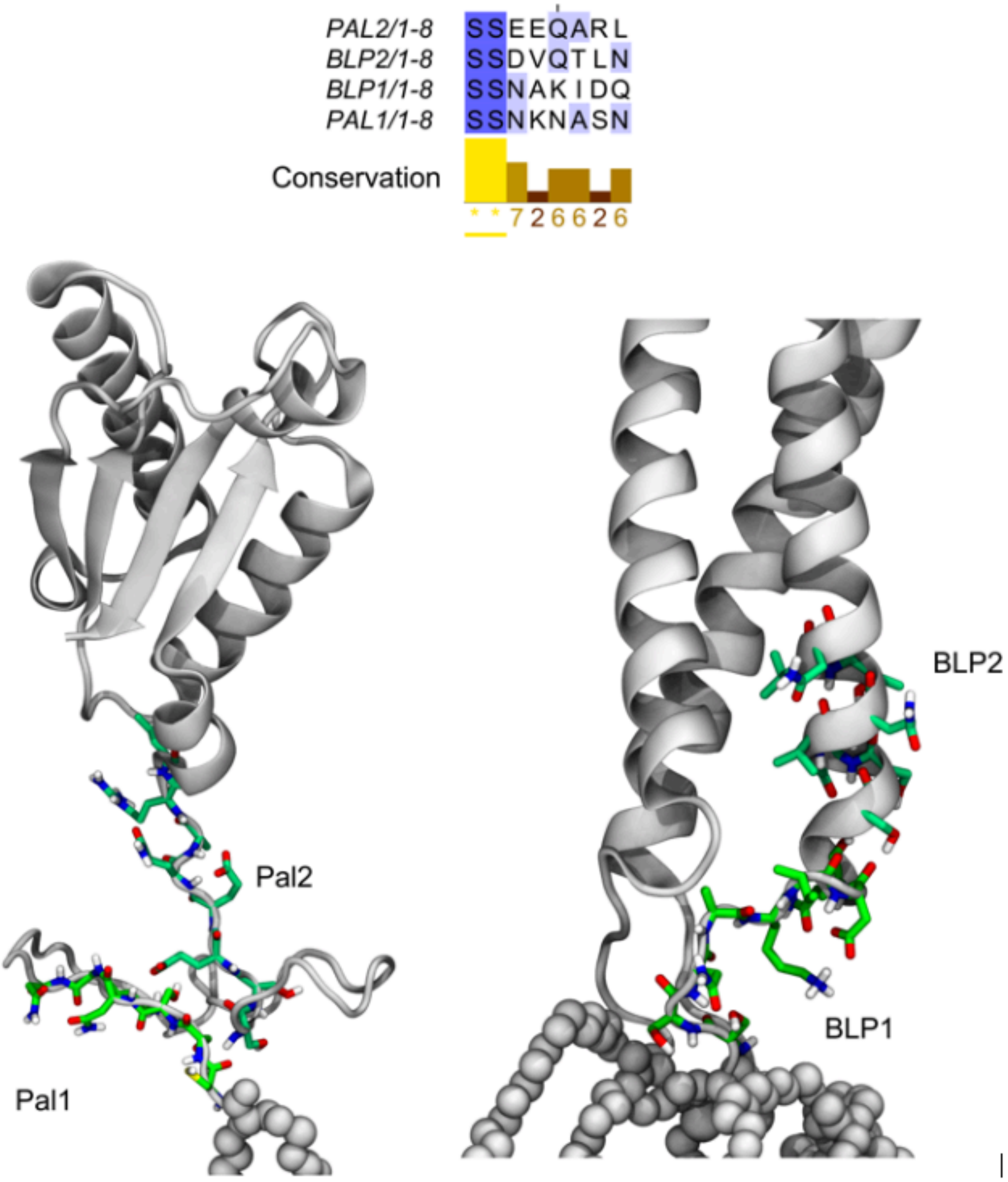


**Figure S9. Short-motifs identified in Pal and BLP proteins. Amino acid sequence alignment for these regions shows the degree of similarity between them. 3D structures show the location of each motif for each protein.**

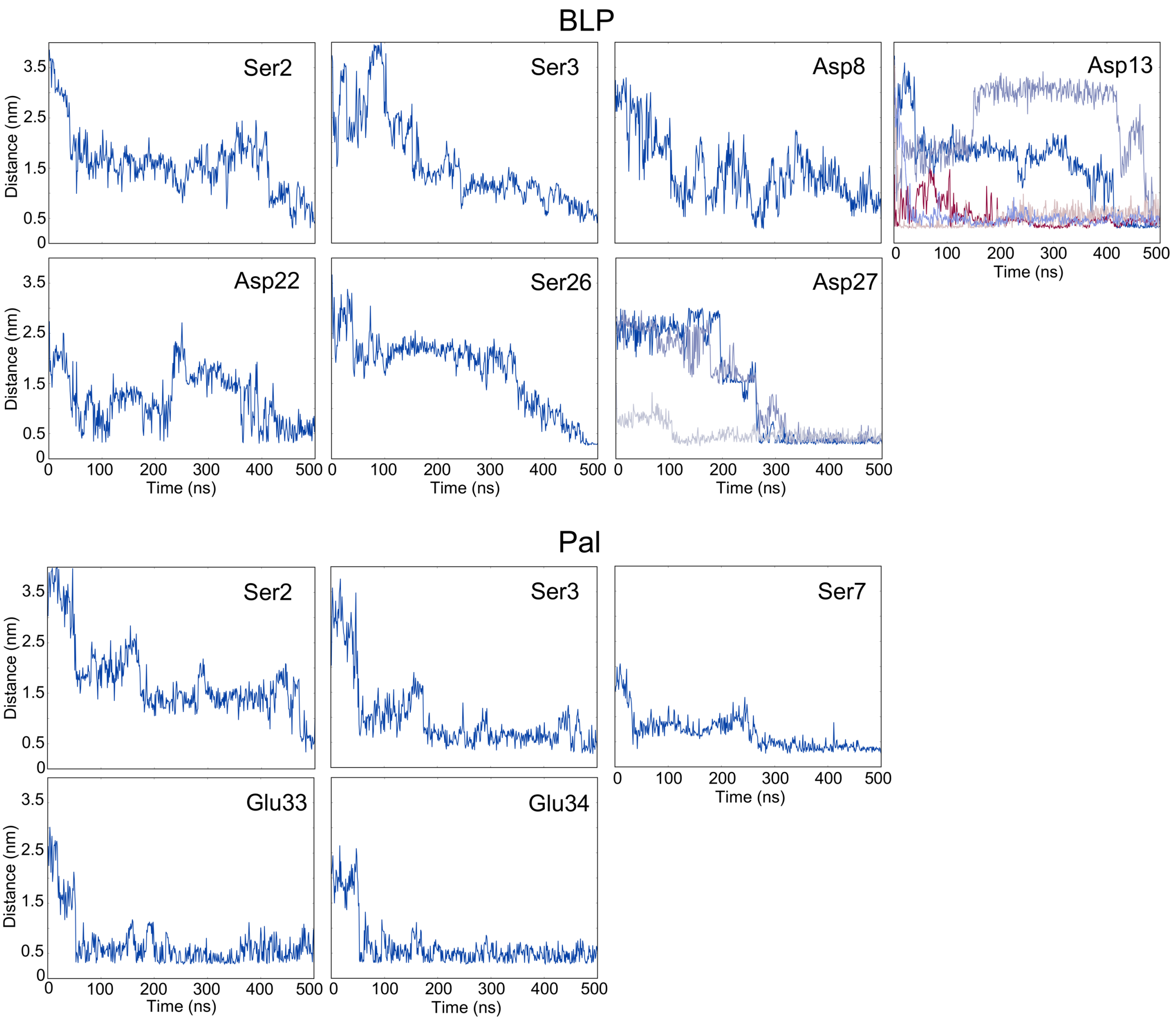


**Figure S10. Distance plots displaying multiple interactions identified during all simulations between PMB1 and Ser, Asp or Glu from BLP and Pal short-motifs. Each curve represents a different interaction between PMB1 and the respective amino acid.**

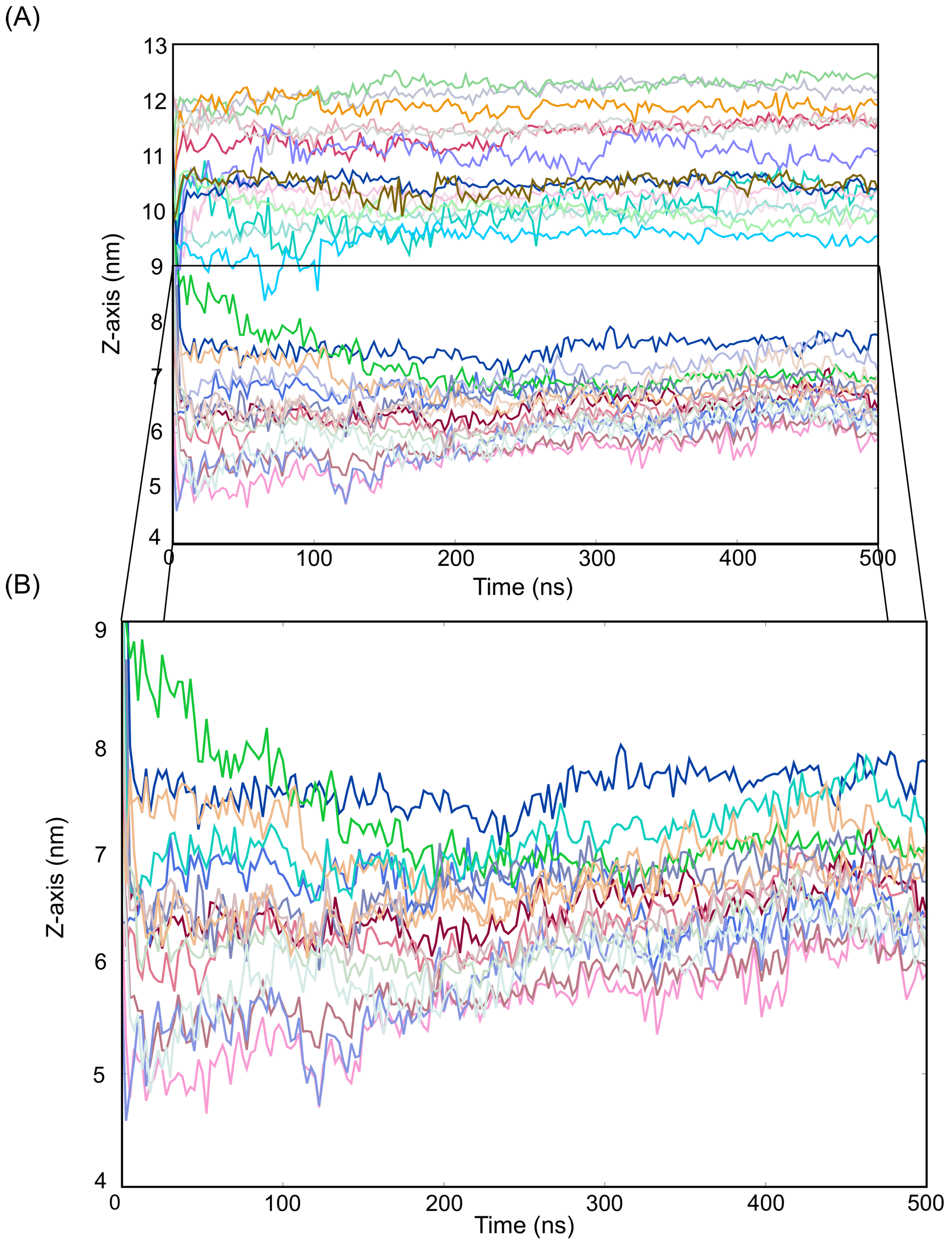


**Figure S11. Z-axis PMB1 motion as a function of time for one PMB1$_{crowd}$ system. Each curve represents a different PMB1 molecule. The zoomed section shows the area where the cell wall is located, especially between 5 and 7 nm.**

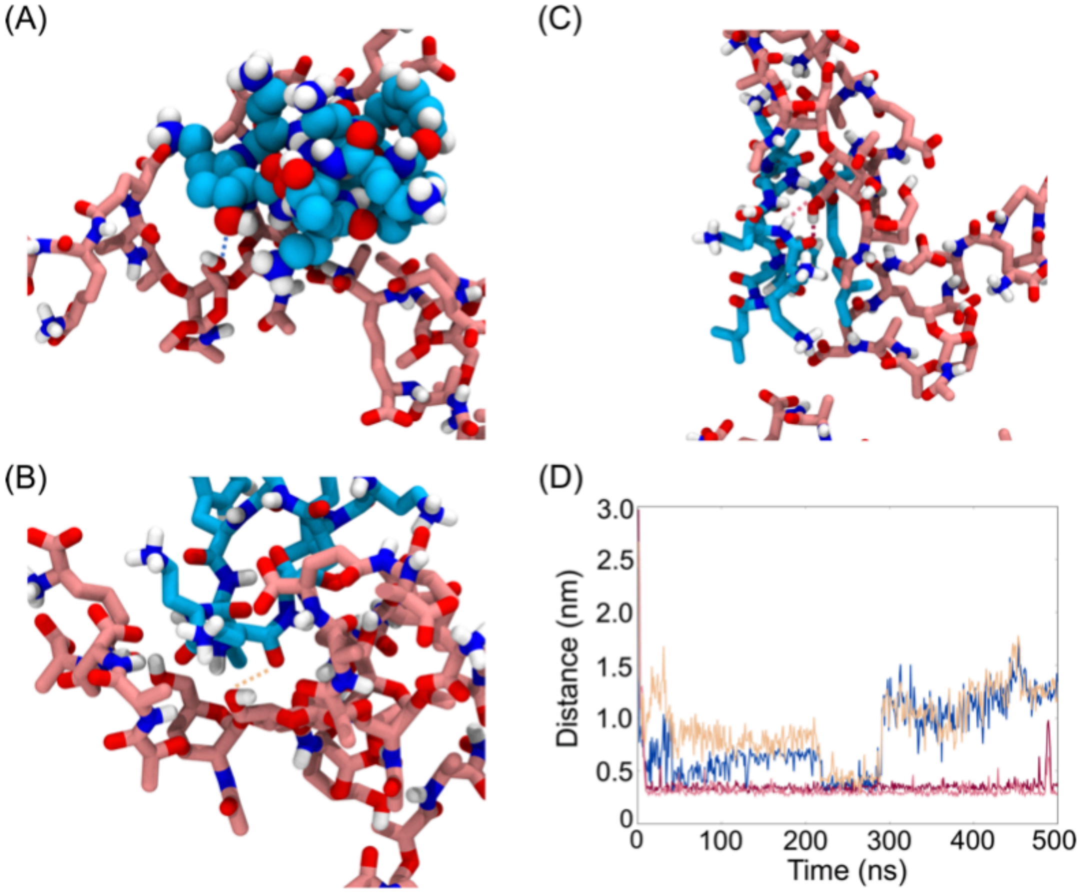


**Figure S12. Hydrogen bond interactions formed between PMB1 molecules and the pyranosidic rings of the cell wall glycan strands. Panels (A-C) depict examples of the different types of interactions observed in the studied systems. The distances for these interactions are plotted as a function of time in panel (D) and colord coded according to dashed lines.**

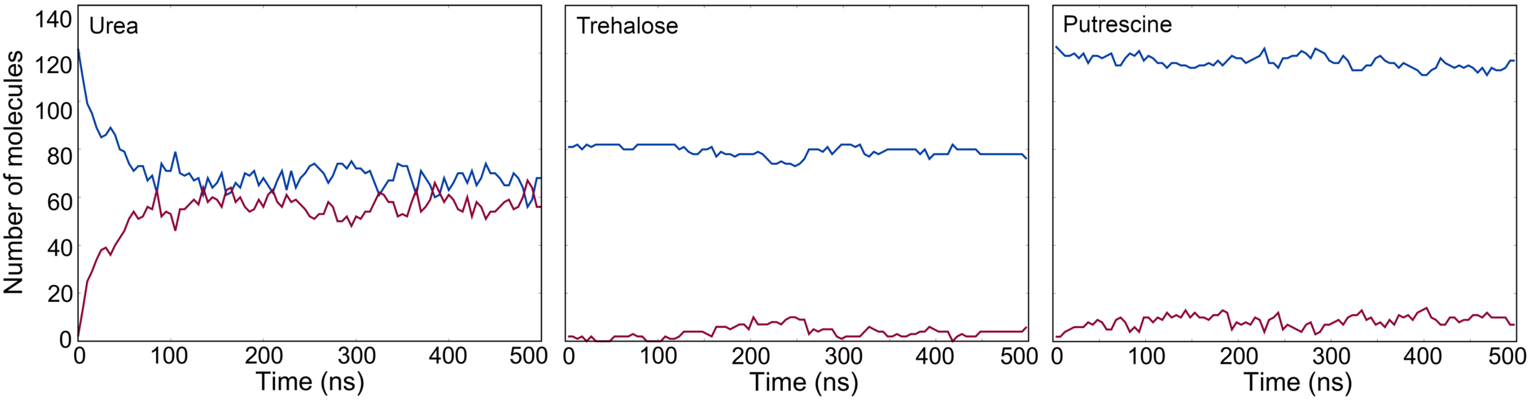


**Figure S13. Number of osmolytes crossing the cell wall as a function of time. Blue curve indicates the number of molecules inside the area delimited by the outer membrane and the cell wall, while the maroon curve shows the number of molecules outside of the mentioned area.**

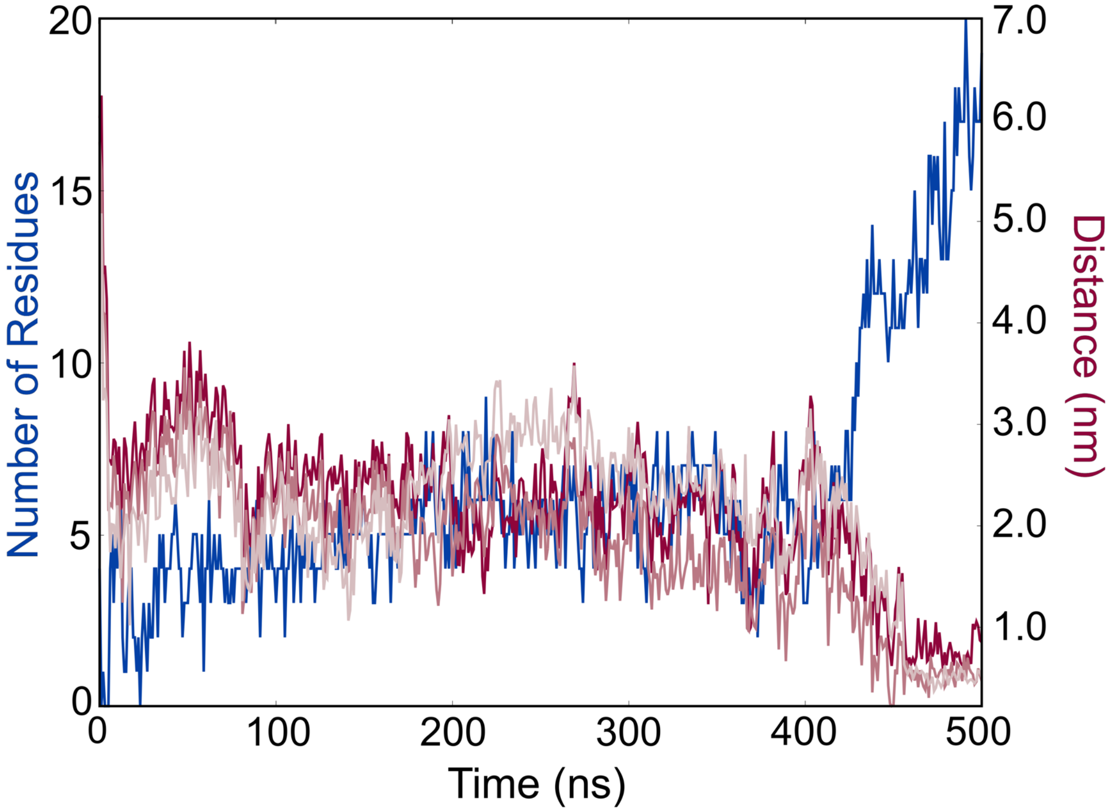


**Figure S14. Number of residues of OmpA that are in close contact (0.4 nm) with the cell wall is shown in blue. The curves with different shades of maroon show the distances between Dab residues of a PMB1 molecule and two DGLu and one m-DAP of the cell wall. The PMB1 molecule is attached to the CTD of OmpA.**

**Table S1.** List of residues interacting with PMB1 in PMBprot and PMBcrowd systems for each protein and description of the binding region where the residues are located.

| System | LolA Residues | LolB Residues | Pal Residues | OmpA Residues | Binding Region(s) |
|---|---|---|---|---|---|
| $PMB_{prot}$ | Phe47, Trp49, Met51, Thr52, Gln53, Pro54, Asp55, Phe72, Glu74 | Phe37, Asp42 Val46, Met100, Ile101, Met107, Ile109, Trp117, Trp148, Gln173 | Ser3, Ser4, Asn5, Lys6, Ser53, Ser54, Glu65, Leu60 Gln63, Gln64, Gln66 | Ala180, Pro181, Ala182, Leu220, Gln223, Leu224, Ser225, Asn226, Asp228, Gly232, Lys230, Ala257, Val260, Val261, Lys267, Gly268, Ile269, Pro270, Ala271, Lys273, Ile274, Ser275 | **LolA**: inside hydrophobic cavity; exterior side near site entrance / **LolB**: hydrophobic site / **Pal**: CTD exterior; in between CTD, linker and membrane / **OmpA**: CTD exterior and near the cell wall; linker region. |
| $PMB_{crowd}$ | Asp31, Met51, Thr52, Gln53, Pro54, Asp55, Ile106, Lys107, Gln108, Phe1113, Lys125 | Ser12, Pro13, Asp14, Trp18, Arg19, Gln22, Pro122, Tyr128 | Ala8, Ser9, Asn10, Gly20, Glu55, Gln53, Gln54, Gln56, Asn93, Arg96, Ser97, Asn90, Pro99, Ser100, Tyr173 | Leu220, Leu224, Ser225, Asn226, Asp228, Gly232, Arg242, Ala247, Lys267, Gly268, Ile269, Ala271, Asp272, Lys273, Ile274 | **LolA**: exterior on both sides near site entrance / **LolB**: N-terminal area / **Pal**: CTD exterior; in between CTD, linker and membrane / **OmpA**: CTD exterior. |
